# Characterization of vertically and cross-species transmitted viruses in the cestode parasite *Schistocephalus solidus*

**DOI:** 10.1101/803247

**Authors:** Megan A Hahn, Karyna Rosario, Pierrick Lucas, Nolwenn M Dheilly

**Author notes:** Address correspondence to Nolwenn M Dheilly.

## Abstract

Parasitic flatworms (Neodermata) represent a public health and economic burden due to associated debilitating diseases and limited therapeutic treatments available. Despite their importance, there is scarce information regarding flatworm-associated microbes. We report the discovery of six RNA viruses in the cestode *Schistocephalus solidus*. None were closely related to classified viruses and they represent new taxa. Mining transcriptomic data revealed the broad distribution of these viruses in Alaskan and European *S. solidus* populations. We demonstrate through *in vitro* culture of *S. solidus* that five of these viruses are vertically transmitted. With experimental infections and field-sampling, we show that one of the viruses is transmitted to parasitized hosts. The impact of these viruses in parasite fitness and pathogenicity, and in host-parasite co-evolutionary dynamics remains to be determined. The detection of six novel viruses in this first characterization of viruses in Neodermatans likely represents a fraction of virus diversity in parasitic flatworms.

## Introduction

Parasitic flatworms (Phylum Platyhelminthes) have long attracted attention for their high prevalence in humans, livestock, and aquaculture animals, and for causing debilitating diseases. Trematodes, commonly known as flukes, and cestodes, known as tapeworms, are of particular interest because around 25-30% of humans alone are currently infected with at least one of these worm species. Several of the pathologies associated with these parasite infections are considered major neglected diseases as they affect countries in the Americas, Asia, and Africa (1). The most notorious example is Schistosomiasis, caused by diverse species of Schistosomes and considered the second most deadly parasitic disease after malaria (2). The blood fluke *Schistosoma haematobium* and the liver flukes *Opisthorchis viverrini* and *Clonorchis sinensis* are recognized as group I carcinogens by the International Agency for Research on Cancer (3–5). Fascioliasis, caused by infection with trematodes from the genus *Fasciola* upon ingestion of contaminated water plants, has detrimental impacts in humans and economically important livestock including pigs, cattle, and sheep (6–8). The most well-known cestodes are *Taenia spp*., *Echinococcus spp.*, and *Hymenolepis nana*. Infections with cestodes are chronic, can remain asymptomatic for long periods, and symptoms are often misdiagnosed, making these diseases difficult to target and treat. Serious effects of parasite infection include cysticercosis and seizures due to *Taenia solium*, cysts or tumors that grow in the liver, lungs, and other organs with Echinococcosis infection, and weakness, headaches, anorexia, abdominal pain, and diarrhea associated with Hymenolepiasis (9–11). Fisheries and, in particular, aquaculture are also largely impacted by parasitic infections due to the high prevalence and densities of worms in fishes that serve as either intermediate or definitive hosts (12).

Despite their high prevalence and negative impacts, diseases associated with parasitic flatworm infections are difficult to prevent or treat. The main method of prevention is avoidance, which can be very difficult in some populations due to limitations in infrastructure and resources (13). Very few pharmaceutical products are currently available for treatment, with Praziquantel and Triclabendazole being the most efficient and commonly used. Moreover, instances of parasite resistance and allergic reactions to these drugs have been reported (14). Thus, researchers have long sought to understand the underlying molecular mechanisms driving host susceptibility and parasite pathogenicity in order to develop alternative therapeutic strategies.

The application of the concept of “holobiont” to parasites, and the recognition that all organisms are associated with microbes suggest that microbes, including viruses, could contribute to parasite pathogenicity (15–18). This realization prompted the launch of the Parasite Microbiome Project (PMP) (16, 18, 19). Viruses of parasitic flatworms remain largely unknown despite the fact that viruses infect all cellular life. The first observations of virus-like particles in trematodes were reported by Jean Lou Justine and its team, studying parasites of mollusks and fishes (20, 21). More recently, Shi et al. (2016) studied the virome of a broad range of invertebrates using metatranscriptomics and reported for the first time the complete genomes of a virus of the order *Bunyavirales* from *Schistosoma japonicum* and of a virus of the family *Nyamiviridae* in the order *Mononegavirales* from a mix of *Taenia sp.*(22). However, no study to date has specifically focused on characterizing viruses of a parasitic flatworm.

As identified by the PMP consortium, there is a need to characterize the virome of parasitic organisms to understand the role of parasites in virus evolution and host-microbe interactions, determine the role of viruses in parasite fitness and host diseases, and identify patterns and processes of host-parasite-virus coevolution (18). We have previously identified *Schistocephalus solidus* as an ideal parasite to answer these questions (15). *S. solidus* is a cestode with a complex life cycle in which the definitive hosts are fish-eating birds and intermediate hosts are a range of cyclopoid copepods and threespine sticklebacks (*Gasterosteus aculeatus*) (23, 24). Since 1946, methods have been developed to culture *S. solidus in-vitro* (25). Eggs can be conserved in the fridge for a few years and hatched to collect coracidia that are used to experimentally infect copepods. Infected copepods are then used to experimentally infect threespine sticklebacks (hereafter ‘stickleback’) (26). This system has been extensively used to identify the cellular and molecular mechanisms involved in host resistance and parasite pathogenicity, and to study host-parasite co-evolution (27, 28). Indeed, these parasites have a broad geographic distribution throughout the Northern hemisphere that parallels the distribution of its highly specific stickleback host. Isolated and genetically distinct populations of fish host and parasite are coevolving within each freshwater lake, providing researchers with an exceptional playground to answer questions that relate to the ecology and evolution of the host-parasite interaction (29). Finally, the genomes of both sticklebacks and *S. solidus* have been sequenced, facilitating the use of molecular approaches (30, 31).

The first step to develop the cestode-host-virus system and facilitate studies on the role of viruses in host-parasite interactions is to characterize viruses associated with *S. solidus*. Herein, we report the discovery of three new species of negative-strand RNA viruses and three new species of double-stranded RNA viruses in *S. solidus*. Mining of Transcriptome Sequence Archives (TSA) and Sequence Read Archives (SRA) data found in GenBank led to the detection of related viral species in *S. solidus* from another continent suggesting that these viruses are widespread. We then tested the prevalence and vertical transmission of identified viruses and evaluated the possibility of cross-species transmission to the hosts through *in vitro* culturing and experimental infections (Figure 1).

**Figure 1:**
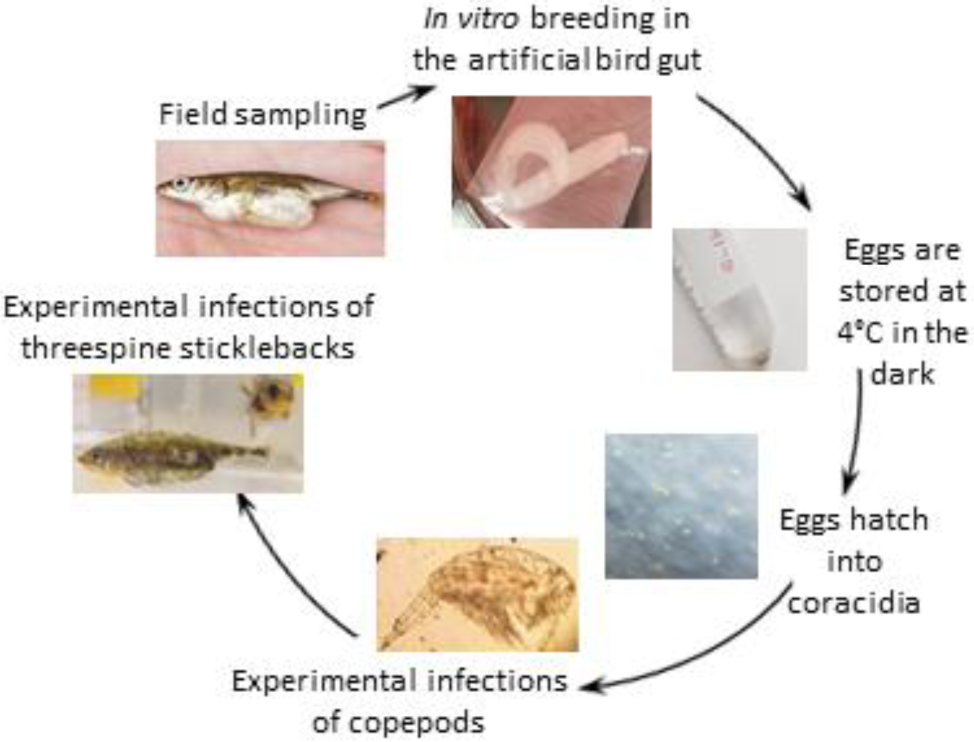
The life cycle of *S. solidus* is reproduced in laboratory conditions to test virus transmission.

## Results

### Virus discovery

To investigate the presence of viruses in the cestode *S. solidus*, viral particles were purified from plerocercoids and coracidia and processed for RNA sequencing. The *de novo* assemblies from two high-throughput sequencing efforts revealed the presence of viral sequences similar to the unassigned Bat rhabdovirus (AIF74284.1), the chuvirus Hubei myriapoda virus 8 (YP_009330113.1), the bunya-like Beihai barnacle virus 5 (APG79235.1), and the toti-like dsRNA viruses Dumyat virus (QAY29251.1) and Hubei toti-like virus 10 (YP_009336493.1). *De novo* assembled contigs were aligned to these reference virus genomes and completed using targeted PCR, RACE and Sanger sequencing as needed. BLAST searches against the genome of *S. solidus* did not yield any matches to these viral sequences, confirming that the identified viruses are not endogenous viral elements (EVE).

The first virus, named Schistocephalus solidus Rhabdovirus (SsRV), contained the five canonical structure domain genes of viruses from the family *Rhabdoviridae*, order *Mononegavirales* and showed a maximum of 59% amino acid (aa) identity to the RdRP of unassigned and partially sequenced Bat Rhabdovirus (AIF4284.1). The SsRV genome encodes for a nucleoprotein (N), polymerase-associated phosphoprotein (P), matrix protein (M), glycoprotein (G) and RNA-dependent RNA polymerase (L) and a short protein in the region between G and L (Figure 1E). All identified open reading frames (ORF) were flanked by conserved transcription initiation (UUGU) and transcription termination/polyadenylation sequences (UC[U]^7^) with very short intergenic region (Table S3). The L protein included common domains such as the *Mononegavirales*-like RdRP domain (pfam 00946), the mononegavirales mRNA capping region V (pfam 14318), a paramyxovirus-like mRNA capping enzyme (TIGR04198) and a *Mononegavirales* virus-capping methyltransferase (pfam 14314).

The second virus, named Schistocephalus solidus Jingchuvirus (SsJV), had a circular genome encoding for a single protein similar to the L protein (Figure 1D) of viruses of the order *Jingchuvirales* with a maximum of 28% aa identity to the RdRP of Hubei Myriapoda virus 8 (YP_009330113.1). The predicted L protein possesses a *Mononegavirales* RdRP domain (pfam 00946), a paramyxovirus mRNA capping enzyme (TIGR04198) and the *Mononegavirales* virus-capping methyltransferase (pfam 14314). No other sequence fragment with similarities to proteins found in viruses of the order *Jingchuvirales* was found.

The third viral genome, named Schistocephalus solidus bunya-like virus (SsBV), had a maximum of 36% identity to the RdRP of the bunya-like virus Beihai barnacle virus 5 (APG79235.1). The longest predicted ORF (Figure 2B) possesses a bunyavirus RdRP domain (pfam04196). Bunyaviruses usually consist of three segments called L, M and S but no other sequence fragment with similarities to viruses of the order *Bunyavirales* was found. Note that SsBV was only found in sequencing data from the total RNA library.

**Figure 2:**
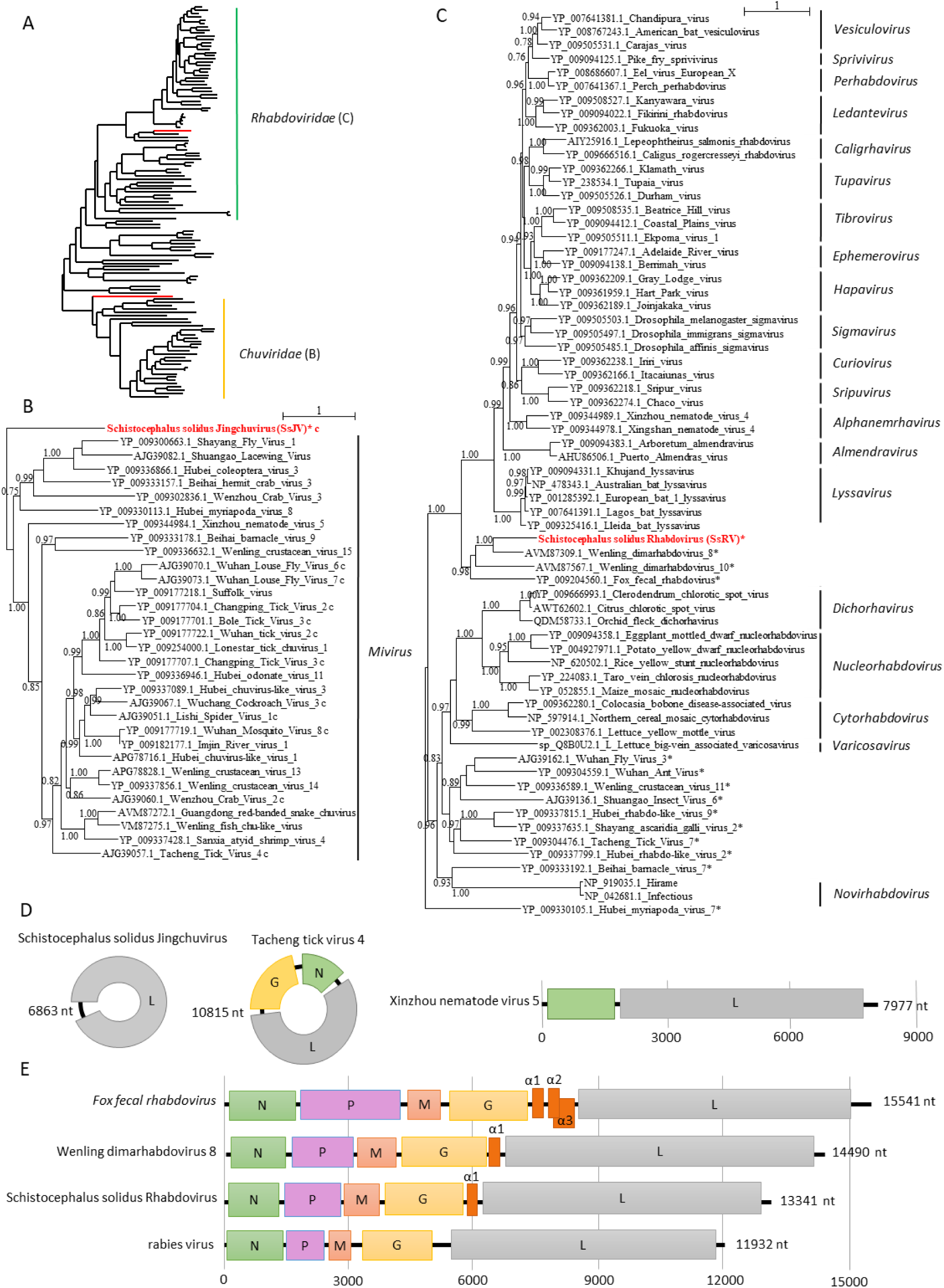
Phylogenetic and genomic characterization of SsRV and SsJV. A) Phylogenetic analysis of the RdRP of viruses from the order *Mononegavirales* and *Jingchuvirales.* The tree was inferred with PhyML using the LG substitution model. B and C) close up view of the phylogenetic tree of the RdRP of viruses from the families *Chuviridae* and *Rhabdoviridae*, respectively. Values next to the branch indicate the results of a Shimodaira-Hasgawa branch test. Genus names and family names are provided next to the branches. * indicates unassigned viruses. c indicates circular genomes. D and E) Genome organization of viruses from *Schistocephalus solidus* aligned to the genome of closely related viruses of the families *Chuviridae* and *Rhabdoviridae,* respectively. Boxes represent putative genes. The black line indicates non-coding regions.

Finally, three sequences similar to viruses of the families *Totiviridae* and *Chrysoviridae* (Figure 3) were found and named Schistocephalus solidus toti-like virus 1 (SsTV1), Schistocephalus solidus toti-like virus 2 (SsTV2), and Schistocephalus solidus toti-like virus 3 (SsTV3). These SsTV viral sequences showed the highest similarities to the partial RdRP sequence of Dumyat virus (QAY29251.1, SsTV1, 27% aa identity and SsTV3, 28% aa identity), or to Hubei toti-like virus 10 (SsTV2, YP_009336493.1, 36% aa identity). All three viruses had two ORFs (Figure 3B), with the second protein encoding for a RdRP similar to Luteovirus, Totivirus and Rotavirus (pfam02123). NCBI Conserved Domain Search (CDD) revealed that SsTV1 ORF1 encodes for a protein with homologies to UL36 large tegument protein of Herpes simplex virus (PHA03247).

**Figure 3:**
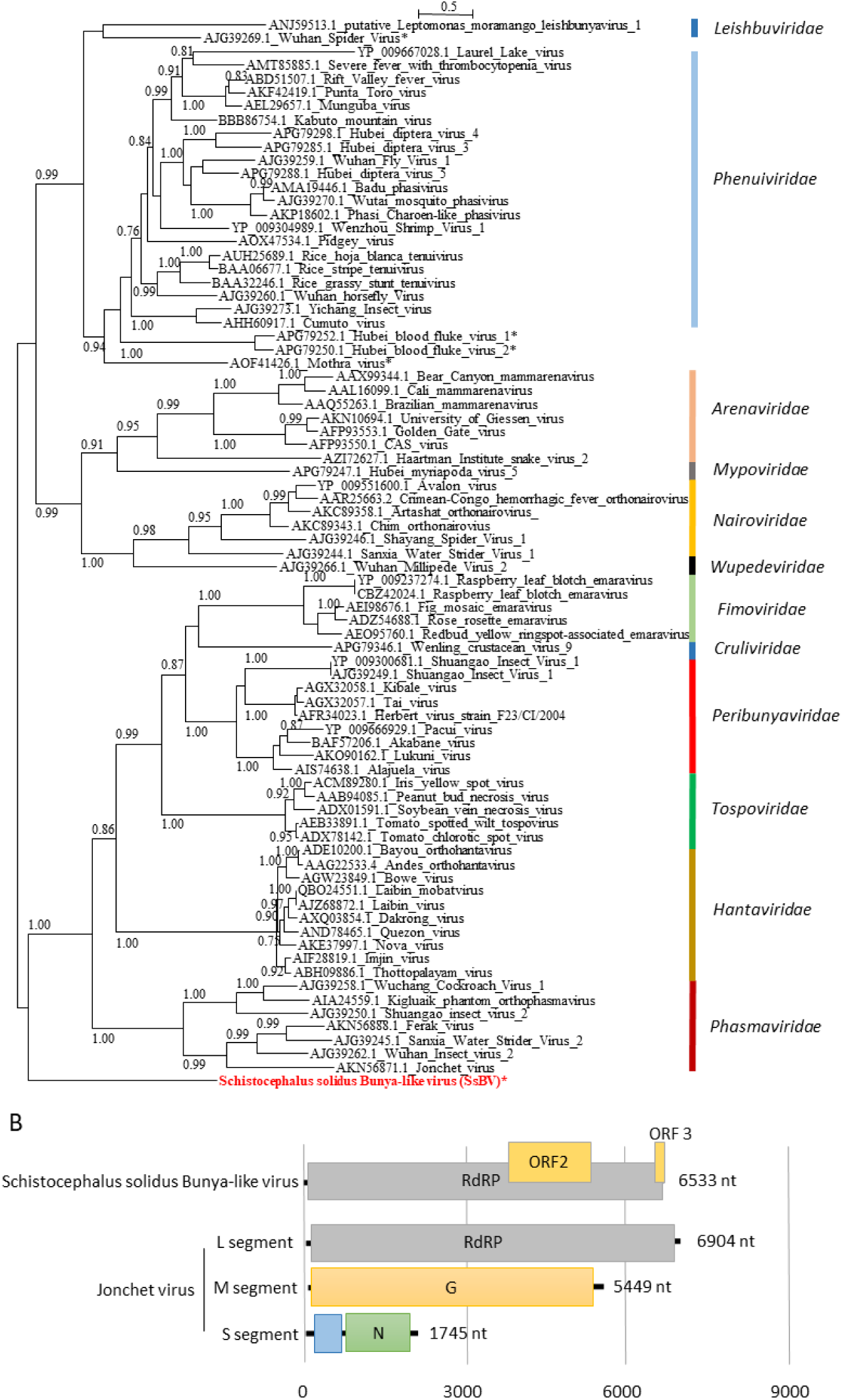
Phylogenetic and genomic characterization of SsBV. A) Phylogenetic analysis of the RdRP of viruses of the order *Bunyavirales*. The tree was inferred with PhyML using the LG substitution model. Values next to the branch indicate the results of a Shimodaira-Hasgawa branch test. * indicates unassigned viruses. Family names are provided next to the branches. B) Genome organization of *Schistocephalus solidus* bunya-like virus aligned to the genome of closely related viruses.

### Evidence for new virus taxa in Schistocephalus solidus

We inferred a phylogenetic tree using the predicted RdRP amino acid sequences from SsRV, SsJV, and 111 representative members of the orders *Jingchuvirales* (31 sequences) and *Mononegavirales* (88 sequences) (Figure 1A). All sequences clustered into previously established genera and families ratified by the International Committee on Taxonomy of viruses (ICTV) except for the *S. solidus*-associated viruses that constituted distinct clades (Figure 1A, Supplementary figure 1). Our results show that SsJV belongs to the order *Jingchuvirales*, but likely represents a distinct taxon from the family *Chuviridae* (Figure 1B). The most closely related viruses, based on the conserved RdRP, is the Hubei myriapoda virus 8 which has a linear genome that encodes for four proteins: a glycoprotein, two hypothetical proteins and the RdRP (Figure S1). The most closely related chuvirus with a circular genome is the Tacheng tick virus 4 that encodes for a glycoprotein, a nucleoprotein and the RdRP.

Our phylogenetic analysis revealed that SsRV belongs to the family *Rhabdoviridae*, grouping closely with Fox fecal rhabdovirus, Wenling dimarhabdovirus 8 and Wenling dimarhabdovirus 10. Notably, SsRV represents a new taxon ancestral to *Lyssavirus* and to the dimarhabdovirus supergroup (Figure 1C). The Bat rhabdovirus, Fox fecal rhabdovirus, and Wenling dimarhabdovirus 8 and 10 from fish were discovered from metatranscriptomic studies and host association had not been confirmed and was challenged by the authors (32, 33). The complete genome of the Fox fecal rhabdovirus, Wenling dimarhabdovirus 8 and SsRV were aligned and compared to the genomes of Rabies virus (Figure 1E). In contrast to the Rabies virus, viruses within these new taxa show variable length and seem to be characterized by the presence of one to three small proteins in the region between G and L.

We inferred a second phylogenetic tree using SsBV and the L segment of 78 representative members of all assigned families within the order *Bunyavirales* (Figure 2). Phylogenetic analysis confirmed that SsBV has no close known relatives and likely constitutes a new family of viruses. SsBV was not found to be closely related to the bunya-like viruses discovered in the trematode *Schistosoma haematobium* (Hubei blood fluke virus 1 and Hubei blood fluke virus 2) (Figure 2A).

We inferred a third phylogenetic tree using SsTV1, SsTV2 and SsTV3 together with 40 viruses representing the families *Totiviridae*, *Chrysoviridae*, and unassigned members closely-related to these families (Figure 3). Phylogenetic analyses of toti-like viruses revealed significant differences between the SsTVs. SsTV1 and SsTV3 cluster together and are most closely related to viruses discovered in other invertebrates including Lophotrochozoa, Nematoda, Crustacea, and Insecta, whereas SsTV2 was most closely related to viruses discovered in insects.

### Mining S. solidus transcriptomic data for viral sequences

At the time of this study, only two transcriptomic studies of *S. solidus* were publicly available. The first study of *S. solidus* transcriptome used 454 GS FLX Titanium sequencing on individuals from Germany and Norway ((34), PRJEB7355, two biosamples). Blast searches against the 454 reads revealed the presence of related strains of SsJV and SsTV1 in one dataset (ERX589070) and of SsTV2 in two datasets (ERX589070 and ERX589072), confirming further the association of these viruses with *S. solidus*.

More recently, the complete transcriptome of *S. solidus* from Clatworthy Reservoir in Somerset, England was assembled using Illumina sequencing and made available as Transcriptome Sequence Archive (TSA; PRJNA304161, 15 individuals) (35). Blast searches against the assembled transcriptome revealed two contigs with high similarity to SsJV, referred to as SsJV2 and SsJV3 thereafter. SsJV2 (GEEE01006270.1) corresponded to the full-length sequence of a variant with 94% amino acid sequence identity to the RdRP of SsJV. SsJV3 (GEE01008921.1) covered only part of the SsJV RdRP, where it shared 63% amino acid sequence identity to SsJV.

We further investigated the presence of viruses within the samples from England by analyzing raw sequencing reads. Reads were assembled for each sample; after removing those that aligned against *S. solidus* genome. Viral contigs were assembled from three of the 15 individuals (SRR2966898, SRR2966894 and SRR2966897): two adult parasites and one mature infective plerocercoid. In addition to the above-mentioned SsJV2 and SsJV3, we assembled a full-length sequence of the L segment of a bunya-like virus, SsBV2, whose full-length genome shares 97.5% aa identity to the RdRP encoded by SsBV. Partial sequences from toti-like viruses similar to SsTV2 were identified in two individuals and were named SsTV4 and SsTV5. The sequences only covered 43% and 31% of the SsTV2 genome length for SsTV4 and SsTV5 respectively. The consensus sequences obtained revealed that both viruses are distinct and display 95% and 52% aa identity to SsTV2. Reads from all 15 individuals were then mapped against these partial genome sequences, revealing that 13 individuals were infected by at least one virus, and many were co-infected by different viruses (Figure S2).

### Prevalence and transmission mode

We tested virus prevalence in plerocercoids from field-sampled sticklebacks from three independent localities in the Matsu Valley, Alaska (Figure 5A, Figures S3-S9). Overall, SsRV was highly prevalent in *S. solidus* in all three tested localities, with an average prevalence of 81% in plerocercoids. In contrast, SsJV, and SsTV2 were detected in 10% and 4% of plerocercoids, respectively, while the remaining viruses (SsTV1, SsTV3, and SsBV) were detected in only 2% of tested plerocercoids. Interestingly, while SsRV, SsJV and SsTV1 were found in all populations, SsTV2 was only found in Cheney and Loberg lakes, and SsTV3 and SsBV were only found in Loberg and Wolf lakes. Only 17.5% of plerocercoids across all populations were free of all viruses.

**Figure 4:**
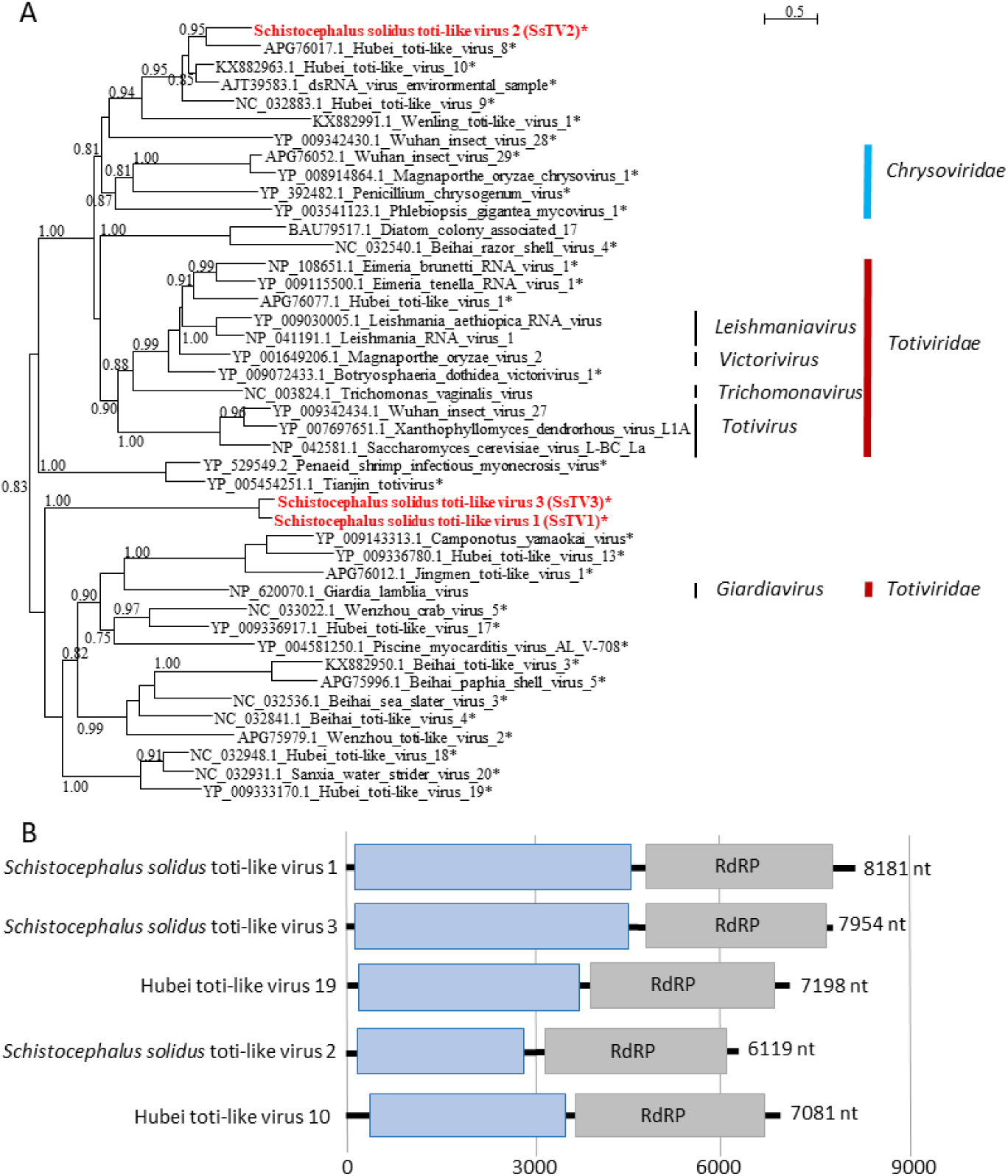
Phylogenetic and genomic characterization of SsTV1, SsTV2 and SsTV3. A) Phylogenetic analysis of the RdRP of toti-like viruses. The tree was inferred with PhyML using the LG substitution model. Values next to the branch indicate the results of a Shimodaira-Hasgawa branch test. * indicates unassigned viruses. Genus names and family names are provided next to the branches. B) Genome organization of toti-like viruses from *Schistocephalus solidus* aligned to the genome of closely related viruses.

**Figure 5:**
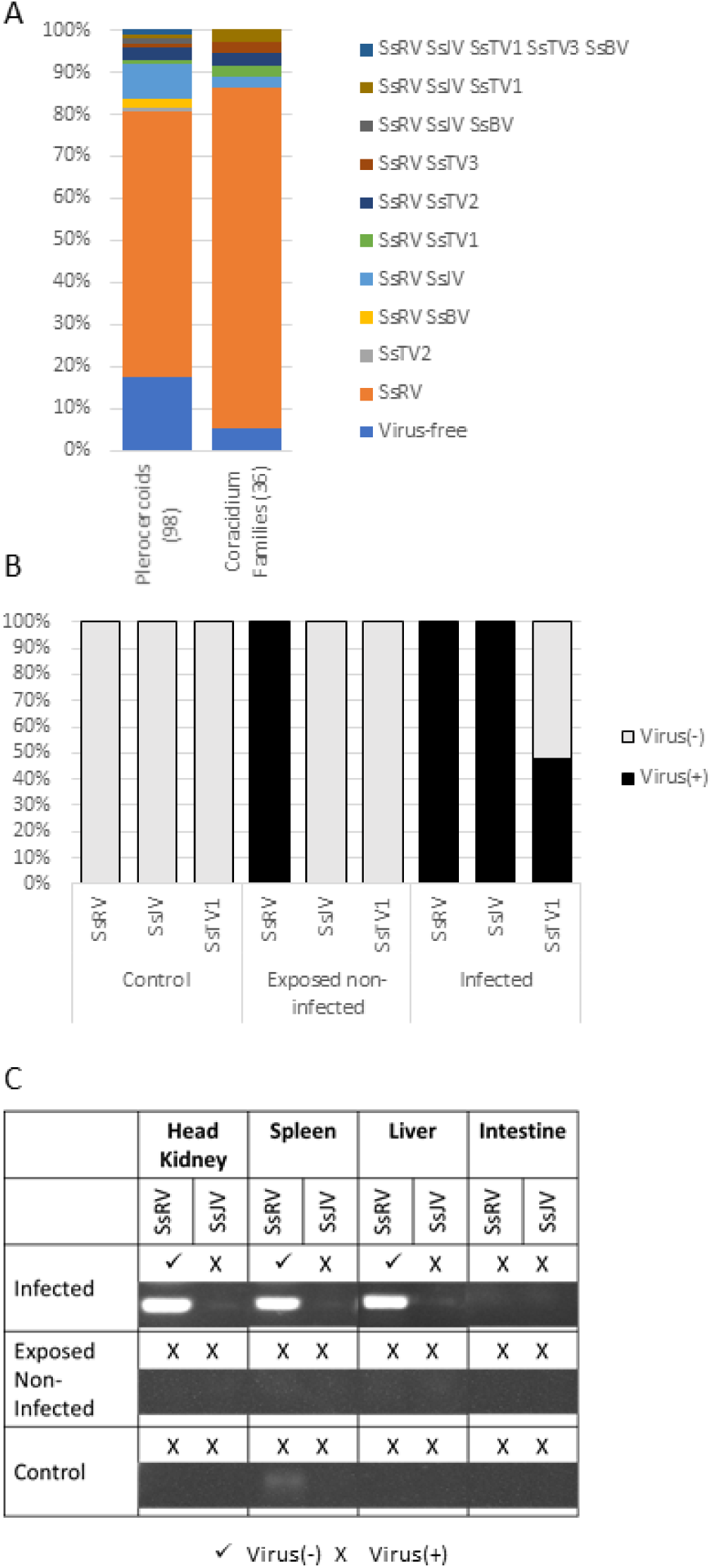
Virus prevalence and transmission over the course of *S. solidus* life cycle. (A) Virus prevalence in plerocercoids from field-collected sticklebacks and in coracidia from *in vitro* generated families. (B) SsRV, SsJV, and SsTV1 presence was assessed in copepods experimentally infected by *S. solidus*. (C) SsRV and SsJV presence was assessed in tissues of sticklebacks experimentally infected by *S. solidus*.

We tested virus presence in worms collected from twenty-four sticklebacks that were co-infected by two or three plerocercoids (Figure 5A, Figures S3 and S10). In many cases, virus-infected and non-infected plerocercoids co-infected the same stickleback host. This was observed for SsRV (eight instances), SsJV (two instances), SsBV (one instance), SsTV1 (two instances), SsTV2 (four instances), and SsTV3 (two instances). Overall, 79% of the plerocercoids from co-infected sticklebacks were SsRV(+), which is not significantly different from the overall prevalence in the populations and suggests little to no horizontal transmission at this developmental stage.

We tested the presence of *S. solidus*-associated viruses in coracidia obtained from *in vitro* breeding to assess the potential for vertical transmission. Both complete genome sequencing and diagnostic PCR results indicated the presence of viruses in lab bred coracidia (Figure 5A). Among the 38 families that were obtained from outbreeding plerocercoids *in vitro*, thirty-four families were SsRV(+), four families were SsJV(+), one family was SsTV1(+), one family was SsTV2(+), and one family was SsTV3 (Figure S9). None of the families were SsBV positive based on PCRs and we later confirmed that none of the plerocercoids randomly selected for breeding were infected by this virus. The lack of lab bred SsBV(+) families prevented further studies and testing of vertical transmission for this virus. The presence of SsRV, SsJV, SsTV1, SsTV2, and SsTV3 in coracidia indicates that these viruses are vertically transmitted.

To further determine the rate of vertical transmission, we experimentally infected copepods with coracidia hatched from virus(+) families. The presence of viruses in procercoids was then assessed for 50 individuals (Figure 5B). While SsTV2 and SsTV3 were found in coracidia, these families showed very low hatching success, preventing us from conducting experimental infections of copepods to test the rate of vertical transmission. SsRV and SsJV were found in all 50 tested procercoids, indicating a 100% success of vertical transmission of both viruses (Figures S11-S13). In contrast, 48% of the procercoids were infected by SsTV1 (Figure S13).

### Cross-species transmission to the hosts

To test SsRV, SsJV, and SsTV1 cross-species transmission to the first intermediate host, cyclopoid copepods, we tested their presence in exposed but non-infected copepods. We did not test their presence in infected copepods as the dissections were too subtle to ensure the absence of contamination from *S. solidus*. SsRV, but not SsJV and SsTV1, was found in exposed but non-infected copepods (Figure S13). No virus was found in control non-exposed copepods. To confirm that no contamination from *S. solidus* was present in exposed but not infected copepods, we used *S. solidus* specific primers to attempt to detect the parasite (Figure S15). *S. solidus* was not found in any of these individuals. However, as we were unable to test if the viruses were associated with copepod tissues or only present on the copepods surface.

To assess the potential for *S. solidus*-associated viruses to be transmitted to the intermediate fish host, we conducted experimental infections of sticklebacks with individual copepods infected with SsRV(+) or SsJV(+) parasites. Since SsTV1 only had a 48% success of transmission it was excluded from this experiment. All four stickleback successfully infected by a SsRV(+) parasite carried the virus within their liver, spleen and head kidney, but the virus was absent from the fish gut (Figure S16). The one fish that was successfully infected by a SsJV(+) parasite did not transmit the virus to its fish host (Figure S16). Exposed but non-infected sticklebacks and control non-exposed sticklebacks were not infected by either virus (Figure 4).

To further test for the rate of cross-species transmission of the highly prevalent SsRV to the fish intermediate host, the presence of SsRV was tested in the liver of field-sampled sticklebacks that were infected by SsRV(+) parasites. Our results showed the presence of SsRV in the liver of all 24 stickleback infected by a SsRV(+) parasite (Figure S18). Among those, 6 sticklebacks were co-infected by both SsRV(+) and SsRV(-) parasites, and yet the virus was found in the fish liver tissue.

To assess the potential for SsRV, SsJV, SsTV1, SsTV2 and SsTV3 to be transmitted to infected definitive hosts, we tested the presence of viruses within the secretory products of breeding adult *S. solidus*. We found SsRV in the culture medium used for breeding all twelve SsRV(+) families whereas we did not find SsJV, SsTV1, SsTV2, or SsTV3 in the culture medium that was used for breeding families (Figure S17).

## Discussion

### A glimpse into a parasitic flatworm viral diversity

In the current study, we used viral purification and shotgun sequencing to identify viruses from the cestode *S. solidus.* We purified viruses from few individuals from three lakes in the Matsu Valley, Alaska, and discovered six new species of viruses; a negative strand RNA virus of the order *Mononegavirales* and family *Rhabdoviridae* (SsRV), a negative-strand RNA virus of the order *Jingchuvirales* (SsJV), a negative sense RNA virus of the order *Bunyavirales* (SsBV) and three double stranded toti-like viruses from unassigned taxonomic groups (SsTV1, SsTV2, and SsTV3). We subsequently found that SsTV1, SsTV2, SsTV3, SsBV, and to a lesser extent SsJV have very low prevalence in field-collected specimens. Genotyping has revealed that *S. solidus* populations remain distinct in individual lakes despite the high motility of its avian definitive host, and different parasite clades co-exist on a given continent (36). Given our sequencing of relatively few individuals in a small number of lakes from a restricted geographic area, it is likely that sequencing purified viruses from a greater number of individuals, and extending the geographic area, will reveal the presence of a greater diversity of *S. solidus*-associated virus species. In addition, *S. solidus* can be found in freshwater lakes throughout the Northern hemisphere (28, 37–39). Mining the *S. solidus* transcriptomic data generated from few individuals from England and Germany, revealed related species of jingchuviruses, bunya-like viruses and toti-like viruses. This confirms that *S. solidus* populations in other lakes and on other continents are infected by different strains of the same viruses reported here. Due to their short generation time, viruses can rapidly diverge when physically isolated in different host populations, increasing viral diversity. Given the great genetic diversity of *S. solidus*, mediated mostly by geographic isolation, but also by selection pressures imposed by the stickleback host (36), *S. solidus* most likely hosts a much greater diversity of viruses than those identified here by sampling a tiny fraction of the parasite’s geographic range and genetic diversity. Further investigations of viruses associated with *S. solidus* could unravel the role of parasite broad geographic distribution and strong genetic structure in virus diversification.

Parasitic flatworms have specialized in parasitizing vertebrates and literally all vertebrates are parasitized by at least one species of parasitic flatworms. They constitute a very diverse and successful phylum. Here, SsBV and SsBV2 were not closely related to recently sequenced bunyaviruses discovered in Schistosomes, who were more closely related to the family *Phenuiviridae* (*22*). This result hints at the potentially large diversity of bunya-like viruses associated with Neodermatans and suggests that a more comprehensive characterization of viruses-associated with parasitic flatworms will likely lead to the discovery of many new viral taxa.

### Genome completeness

While the full-length genomes of SsRV, SsTV1, SsTV2, and SsTV3 were obtained, it is unclear whether SsJV and SsBV genomes are complete. Our assemblies of SsJV and SsBV revealed a single genome that encodes for the RdRP. SsJV belongs to the recently accepted order *Jingchuvirales* within the class *Monjiviricetes*. The order currently contains only the family *Chuviridae* and the genus *Mivirus* with 29 species (40). Viruses within this order have small, often segmented and circular genomes (41). Similarly, SsBV belongs to the order *Bunyavirales* that includes viruses with segmented genomes consisting of two to six fragments that are packaged stochastically (42). We were unable to find any other sequence fragments related to chuviruses or bunya-like viruses in our samples or in the transcriptome of *Schistocephalus solidus* probably because additional SsJV and SsBV segments are significantly divergent from known viruses, hindering our ability to detect these segments through sequence similarity-based searches. Another possibility is that SsJV may be satellite or helper virus encapsidated by another virus (43). This could explain the fact that SsJV was detected in samples after nuclease treatment, its high rate of vertical transmission and that it was never found in an individual that was not already infected by SsRV. SsBV, however, was absent from sequencing purified viruses and was only discovered because we sequenced total RNA, which may indicate its sensitivity to our viral purification strategy (e.g., nuclease treatments). Thus, an alternative hypothesis is that SsBV lacks capsids or envelop proteins and relies solely on vertical transmission similarly to viruses of the family *Narnaviridae*.

### Phylogenetic position

Our phylogenetic analyses revealed that all newly discovered viruses are distinct from the known diversity of viruses and constitute new undescribed taxa. These viruses also often had an ancestral position to other viruses suggesting that viruses of parasitic flatworms may have played a role in virus evolution. Previous studies showed that the order *Jingchuvirales* has an ancestral position to the order *Mononegavirales* (41). SsJV appears to have an ancestral position to other viruses within the order *Jingchuvirales* and hence, may be among the most ancestral of all known viruses within the orders *Mononegavirales* and *Jingchuvirales*. Similarly, SsTV1and SsTV3 clustered together and had an ancestral position to all closely related viruses found in other invertebrates. SsBV did not cluster with known viruses and its phylogenetic position fell between the cluster of viruses of the families *Phasmaviridae*, *Hantaviridae*, *Tospoviridae*, *Peribunyabiridae*, *Cruliviridae* and *Fimoviridae*, and the cluster of viruses of the families *Phenuiviridae*, *Arenaviridae*, *Mypoviridae*, *Nairoviridae* and *Wupedeviridae*, again indicating an ancestral nature to many other viruses. A more robust interpretation can be made from the phylogenetic position of SsRV that suggests an ancestral role of parasitic flatworms in the evolution of the *Lyssavirus*. The family *Rhabdoviridae* includes viruses of vertebrates, invertebrates, and plants grouped within 20 genera (44, 45). Well known viruses within this family includes the rabies virus, vesiculoviruses and potato yellow dwarf virus that are of public health, veterinary and agricultural importance (46). We found that SsRV is transmitted to *S. solidus* intermediate copepods and stickleback hosts over the course of infection and could potentially be transmitted to its definitive avian host during breeding, which would facilitate host switch from a parasitic flatworm to its vertebrate host. Based on SsRV phylogenetic position, and with the support of these experimental results, we propose the following evolutionary scenario for the dimarhabdovirus supergroup: An ancestral rhabdovirus of a parasitic flatworm acquired the ability to replicate exclusively in vertebrates, diverging to an ancestral Lyssavirus. Then, an ancestral virus of the dimarhabodvirus supergroup acquired the ability to use insects as vectors to increase its transmission among vertebrate hosts (47). Finally, rare additional host shifts events can explain the host association of different genera within the dimarhabdovirus supergroup (47). Parasites have a close and intimate relationship with their hosts that could favor virus host shifts. Multi-host parasites, such as parasitic flatworms, have the potential to acquire or transmit viruses from and to each of their hosts, thus providing the means for viruses to complete major host shifts across distantly related host taxa. Clearly, future studies characterizing a greater diversity of viruses of parasitic flatworms and their phylogenetic position relative to the diversity of viruses within their intermediate and definitive hosts has the potential to fill major gaps in our understanding of virus evolution.

### Transmission and impact on parasite fitness

Viral infections may have implications for the ecology of their parasitic hosts. Viruses of parasites can either be beneficial for the parasite by increasing infectivity or transmission to the next host, or they can be hyperparasitic and result in parasite hypovirulence that benefits the parasitized host (17). The nature of virus interaction with its parasitic host will have downstream effects on the interaction between host and parasite and on the evolution of the combined system. Indeed, hyper- and hypovirulent viruses would displace the parasite virulence level away from the optimal evolutionary strategy (ESS) and act as selection pressures on virulence evolution (48). ESS theory predicts that strict vertical transmission should be rare excepting if the virus provides fitness advantages (49). However, we found five viruses (SsRV, SsJV, SsTV1, SsTV2, and SsTV3) that are vertically transmitted. The one virus that was not found in coracidia, SsBV, was not present in any parasite used for breeding preventing us from testing the vertical transmission of SsBV, which remains unknown. Our experimental design did not allow us to determine whether only one or both parents were infected by the tested viruses at the time of breeding because eggs remained attached to the surface of adult worms after breeding. Therefore, it is unclear whether the 48% rate of vertical transmission of SsTV1 is due to only one parent being infected by this t. Similarly, the 100% rate of vertical transmission observed with SsRV(+) and SsJV(+) could result from both parents being infected by the virus and vertical transmission of viruses by mothers only (50). In the future, this experimental design could be improved by collecting small tissue samples of adult worms before breeding in order to assess the viruses’ presence in each parent. Regardless of the limitation of our design, the high rate of vertical transmission suggests that SsRV, SsJV, and SsTV1 probably have low virulence, or may even be beneficial for *S. solidus*. This is further supported by the presence of virus(+) and virus(-) parasites in co-infected sticklebacks indicating that these viruses have a low rate of horizontal transmission at this developmental stage.

SsRV maintained high prevalence in all three tested populations and is cross-species transmitted to its hosts, further suggesting that it could be beneficial for the parasite or detrimental to the parasitized hosts. The absence of SsRV in the stickleback intestine indicates that the virus is likely transmitted to the host while the parasite is developing to sexual maturity in the body cavity (51). SsRV was found in the muscle of the body cavity, and in the spleen, liver and head kidney of parasitized sticklebacks. The fish liver, spleen and head kidneys are involved in many biological processes in sticklebacks, such as immune response to infection by *S. solidus*, metabolism and energy storage (52–58). The ability of the virus to replicate in stickleback cells, and its impact on host immune response to parasite infection remains to be assessed but its presence is likely sufficient to stimulate the host immune system. For example, SsRV could be used as a biological weapon to deter the host immune response away from the parasite (17). Other studies have revealed the presence of RNA viruses in the human parasites *Trichomonas vaginalis*, *Giardia lamblia,* and *Leishmania spp.* (59–62). These viruses impact interactions between host and parasite: the *Trichomonavirus,* and *Leishmaniavirus* exacerbate virulence of their parasitic hosts whereas *Giardiavirus* is associated with a decrease in parasite virulence (63). The *Trichomonavirus* and *Leishmaniavirus* induce Type I interferon (IFN) and elevated proinflammatory response that controls the severity of the diseases (64, 65). Examples of parasitic flatworms infection that impede antiviral immunity and are associated with increased vial load are abundant, but in some cases, a protective effect has also been observed (66, 67). The transmission of viruses of parasitic flatworm to hosts can explain apparent cross-reactions of the immune system. Interestingly, even though SsRV was not found in any fish tissues when parasite infection was not successful, it was found in copepods that were exposed to *S. solidus* but resisted infection. We used *S. solidus* specific primers to confirm that the parasite was no longer present in exposed copepods and ensured that the virus was indeed associated with the copepod. It remains to be determined if SsRV can replicate in copepods and infect *S. solidus* that the copepod may encounter at a later time. Copepods could potentially serve as vectors or reservoirs of SsRV, facilitating horizontal transmission between procercoids. That being said, our knowledge of *S. solidus* prevalence and exposure rate in copepods in field settings is very limited. Therefore, we can only speculate regarding the potential role of copepods in SsRV ecology.

In contrast, SsJV, SsTV1, SsTV2, SsTV3, and SsBV had low prevalence in all Alaskan tested populations. The low prevalence of SsJV and SsTV1 is particularly surprising given the estimated high success of vertical transmission. This result suggests that at some point during the parasite life cycle, SsJV(+) and SsTV1(+) parasites are less successful than SsJV(-) and SsTV1(-) parasites and that these viruses may negatively impact *S. solidus* fitness. Alternatively, the low prevalence of SsJV and SsTV1 may result from competition with the highly prevalent SsRV or a lower success rate of vertical transmission from individuals co-infected with SsRV. In support of this second hypothesis, SsRV was not found in transcriptomic data of S. solidus from Europe whereas Jingchuviruses and Toti-like viruses were present. Given the very low number of individual worms (two and fifteen respectively for each Bioproject), the discovery of these viruses suggests that their prevalence might be higher in Europe. Functional experiments to determine the fitness impact of these viruses, alone and in combination, on *S. solidus* and its stickleback host need to be conducted.

### Perspectives

Viruses and parasites alike have significant impacts in health sciences, but until recently they have mostly been studied separately by virologists and parasitologists, respectively. Herein, we discovered vertically transmitted viruses in the cestode *S. solidus*, and showed that at least one of these can be transmitted to parasitized hosts. Given the importance of viruses and their prevalence in all cellular organisms, any parasitic flatworm should be considered a holobiont and the presence of associated microbes should be investigated (16). The viruses we discovered in *S. solidus* can serve as reference to facilitate the search for related species in other parasitic flatworms. For parasitologists, it is like a Pandora’s box has been opened, with a myriad of new possibilities for understanding parasite-associated diseases, and for the development of therapeutic strategies. Characterizing the role of parasite viruses in host-parasite interactions will allow us to identify the real culprits for observed symptoms, or diseases, a necessary step towards the development of new targeted therapies to treat or prevent debilitating diseases that have been plaguing populations for decades. A striking example of the therapeutic potential is the successful development of a vaccine that target the *Leishmaniavirus* and provide a cell-mediated immune protection that has the potential to reduce exacerbated forms of leishmaniasis (68).

## Materials and Methods

### Initial sample processing and sequencing for virus detection

*Schistocephalus solidus* field-collected specimens were initially screened for the presence of viruses through viral purification and shotgun sequencing. For this purpose, *S. solidus* plerocercoids were dissected out of four sticklebacks collected in Cheney Lake, Alaska (61° 12’ 17” N, 149° 45’ 33”) in June 2016 resulting in four parasite samples. The plerocercoids were cut into pieces and immediately transferred into phosphate-buffered saline (PBS) for virus purification through filtration followed by chloroform and nuclease treatment according to Ng et al (69, 70) with some modifications to the protocol. Briefly, tissue samples were homogenized in sterile PBS by bead beating with 3 mm glass beads. The homogenates were centrifuged for 1 min at 6000 rpm and pellets were discarded. The recovered supernatants were further diluted with 500 µl of PBS and centrifuged at 6000 rpm for 6 min to remove remaining cell debris. The supernatants were then filtered successively through 0.4 µm and 0.22 µm sterile cellulose acetate filters (Corning CAT# 8162) and filtrates containing the viral fraction were incubated for 10 min in 0.2 volumes of chloroform. The viral fraction was then recovered from the aqueous phase after centrifugation for 20 seconds at 20,000 rpm. A second chloroform treatment was applied to ensure removal of bacterial contaminants. The viral fraction was further purified by treating with 2.5 U of DNase I and 0.25 U of RNase A at 37°C for 3 hours to eliminate non-encapsidated DNA and RNA. EDTA (pH = 8, Sigma Aldrich CAT# E7889) was added at a final concentration of 20mM to inactive nucleases prior to nucleic acid extraction.

Viral DNA and RNA were simultaneously extracted using the QIAamp Mini Elute Virus Spin Kit according to manufacturer’s instructions. DNA was then removed using a Turbo DNAse treatment (Thermofisher CAT# AM1907). Four sequencing libraries, one per parasite sample, were prepared with the NuGen Oviation Universal RNA-Seq System (CAT#0343) following the standard protocol and 18 PCR cycles. Libraries were used for single-end sequencing (1 × 150bp) on an Illumina Hi-Seq 4000 (Institute of Biotechnology at Cornell University). We obtained 10.22, 12.57, 10.81 and 1.54 million reads for each respective sample. Sequences were processed through Stony Brook University Seawulf high performance computing cluster. For each dataset, adapters were removed using Trimmomatic version 0.36 with default settings and PhiX174 contaminants were removed using Bowtie 2 (--very-sensitive-local) (71). Sequence quality after trimming was verified with FastQC version 0.11.5 (72). D*e novo* assembly was completed by pooling sequence data from all four samples using Trinity (31). Contigs representing partial viral sequences similar to various rhabdoviruses and chuviruses were identified through BLAST searches against GenBank non-redundant database (BLASTx, e-value < 10^−10^). The partial sequences represented two viruses, a rhabdovirus, named Schistocephalus solidus Rhabdovirus (SsRV) and a chuvirus named Schistocephalus solidus Jingchuvirus (SsJV).

### Sampling and *in-vitro* culturing of *S. solidus* for virus genome sequencing and experimental infections

#### Field sampling

The detection of viral sequences in plerocercoids collected from Cheney Lake prompted further sampling of *S. solidus* from various lakes to complete detected viral genomes, screen for other viruses, evaluate the prevalence and distribution of detected viruses, and perform experimental infections (Figure 1). In June of 2018, 31, 20, and 46 plerocercoids were collected from sticklebacks fished from Cheney Lake, Wolf Lake (61° 38’ 36” N, 149° 16’ 32” W), and Loberg Lake (61° 33’ 33.5” N 149° 15’ 28.9” W), respectively. For a subset of stickleback hosts, the liver was collected. Whole plerocercoids and fish tissue samples were transferred into RNA later for future analyses.

#### In-vitro culture of S. solidus plerocercoid*s*

Freshly collected plerocercoids were used for *in vitro* breeding by placing size-matched pairs into sealed biopsy bags (26, 50, 73). Each pair was incubated for 4 days at 40°C into 250ml of Minimum Essential medium (MEM Sigma M2279) enriched with HEPES buffer (Sigma CAT# 83264, 50ml l^−1^), Antibiotic antimycotic (Sigma CAT# A5955, 10ml l^−1^), L-glutamin (Sigma CAT# G7513, 10mmol l^−1^) and glucose (Sigma CAT# G7021, 40ml l^−1^). Eggs were collected and the culture medium was replaced every 48 hours for 4 days. Upon collection, the eggs were washed 5 times in sterile water and stored at 4°C in the dark. Families were bred for parasites from each lake, resulting in 16 families from Cheney lake, 13 families from Wolf lake, and 9 families from Loberg lake. Plerocercoids used for breeding were then transferred into RNA later and a sample of culture medium was mixed V/V with RNA later for future analyses.

#### Egg hatching and processing for virus detection and sequencing

Newly hatched coracidia from *S. solidus* families from Wolf Lake (six families), Loberg Lake (two families), and Cheney Lake (two families) known to carry SsRV or SsJV viruses (detected via PCR, see below) were used for a second sequencing effort to complete the genomes and potentially detect more viruses. To stimulate egg hatching, eggs were incubated in deionized water for 3 weeks at 22°C in the dark before being exposed to UV light for 1 hour, placed in the dark overnight and exposed to UV light for 3 more hours (50). Newly hatched coracidia were collected through centrifugation at 6,500 rpm or 5 min at 4°C. To purify viruses, coracidia samples were homogenized in sterile suspension medium (SM) buffer [100 mM NaCl, 8 mM MgSO_4_·7H2O, 50 mM Tris-Cl (pH = 7.5)] through bead beating in a Fisherbrand Bead Mill 4 homogenizer (Fisher Scientific CAT# 15-340-164) for 1 min using a mixture of 0.1 mm and 1 mm glass beads. Homogenates were then centrifuged at 8,000 x g for 10 min and the supernatants containing the viral fraction were filtered through a 0.45 μm Sterivex filter (Fischer Scientific CAT# SVHV010RS) to remove cells. Free DNA and RNA were removed from the viral fraction by incubating filtrates with a nuclease cocktail consisting of 1X Turbo DNase Buffer, 21U of Turbo DNase (Fisher Scientific CAT# AM2238), 4.5U of Baseline-ZERO DNase (Epicenter CAT# DB0711K), 112.5U Benzonase (Fisher Scientific CAT# 707463), and 10 μg/mL RNase A (Fisher Scientific CAT# AM2294) at 37 °C for 2 h. Nucleases were inactivated with 20 mM EDTA prior to nucleic acid extraction.

Viral RNA was extracted from 200 μl of purified viral fraction using the RNeasy kit (Qiagen CAT# 74104) with the on-column DNase digestion step following manufacturer’s recommendations. In addition, total RNA extracts obtained from lab raised plerocercoids and coracidia from each lake (see below) were processed alongside RNA extracts from the purified viral fraction. RNA was reverse-transcribed using the SuperScript IV First Strand Synthesis System (Fisher Scientific CAT#18091050) with random hexamers followed by second-strand cDNA synthesis using the Klenow Fragment DNA polymerase (New England Biolabs CAT#M0212S). The resulting products were cleaned using the AMPure XP Purification system (Beckman Coulter CAT# A63880). Purified cDNA samples from the viral fraction (V) and those from total RNA (T) were pooled into two samples, namely the V-pool and the T-pool. Both pools were fragmented to 300 bp using a Covaris M220 instrument at the Molecular Genomics Core at the H. Lee Moffitt Cancer Center & Research Institute. Next-generation sequencing library construction was performed with the Accel-NGS 1S Plus DNA Library Kit for Illumina Platforms (Swift Biosciences CAT# 10024) following manufacturer’s instructions for DNA inputs <1 ng/μl and 18 cycles of dual indexing PCR for the V-library. For the T-library, fragmented RNA was processed following the protocol for DNA inputs > 10 ng/ul and 10 cycles of dual indexing PCR. Both libraries were commercially paired-end sequenced (2 × 150 bp) on an Illumina HiSeq 4000 System at GENEWIZ.

Sequences were processed through the University of South Florida high performance computing cluster. Raw sequences were trimmed for quality and to remove indexing adapters using Trimmomatic version 0.36.0 (74) with default parameters except for a read head crop of 10 bp instead of zero. Sequence quality after trimming was verified with FastQC version 0.11.5 (72. Due to the high number of indexing PCR cycles, quality-filtered sequences from the V-library were assembled following a pipeline for PCR amplified libraries (75). To do this, sequences were dereplicated using the Clumpify tool from the BBtools package (sourceforge.net/projects/bbmap/). Dereplicated sequences were then assembled using single cell SPAdes (76). Quality-filtered sequences from the T-library were assembled with RNAspades. Contigs larger than 1000 bp were compared (BLASTx, e-value < 10^−10^) against a viral protein database containing sequences from the NCBI Reference Sequence (RefSeq) database (RefSeq Release number 93, https://www.ncbi.nlm.nih.gov/refseq/). This sequencing effort resulted in the detection of contig sequences representing SsJV, SsRV and two novel toti-like viruses, named SsTV1 and SsTV2. As part of an iterative approach, contig sequences were compared against viral sequences detected from parasite datasets, including newly detected *S. solidus* viruses, leading to the detection of a novel bunya-like virus and a third toti-like virus, named SsBV and SsTV3, respectively.

#### Viral genome completion

Quality filtered reads and contig sequences associated with each of the viruses were retrieved by comparing sequences through BLASTn to a database containing newly identified contig sequences, including those from the original assembly done in Trinity, and closely-related sequences. All reads and contigs were re-assembled using the default overlap-consensus algorithm implemented in Geneious version R7. All assemblies resulted in near-complete genome sequences represented by a single contig, with the exception of SsRV for which genome gaps were closed through targeted PCR (primers listed on Table S1). To complete genomes, RNA samples from families originally pooled by lake were screened for each of the viruses to identify positive samples. Positive samples were then used for PCR and rapid amplification of complimentary ends (RACE) assays (77). All PCRs were performed using the AmpliTaq Gold™ 360 Master Mix with GC enhancer (Thermo Fisher Scientific). PCR using primers designed off of the SsJV contig sequence ends confirmed the circular topology of this new chuvirus-like genome (Table S1). The genome ends of the remaining viral sequences were completed through RACE (primers provided in Table S2). Prior to 3’RACE reactions, RNA extracts were denatured at 95°C for 6 minutes and placed on ice immediately to prevent RNA reannealing (78). Denatured RNA was used as template for poly(A) tail reactions using a Poly(A) Polymerase from *E. coli,* which synthetizes poly(A) tails at the 3’ termini of ssRNA templates. Poly(A) reactions contained 1mM ATP, 1X poly(A) polymerase buffer, 0.25 U poly(A) polymerase (New England Biolabs) and 15 ul of RNA. Poly(A)-tailed RNAs were used as template for 3’RACE reactions. The 5’ ends were completed with the 5’RACE system. All RACE products were cloned using the CloneJET PCR Cloning Kit (Thermo Fisher Scientific) and Sanger sequenced using vector primers. All PCR and cloned RACE products were commercially sequenced by TACGen.

#### Egg hatching for experimental infections and diagnostic PCR assays

*S. solidus* eggs were hatched following a similar strategy as that outlined above. Briefly, eggs were allowed to develop in sterile filtered tap water for 3 weeks at 18°C in the dark. Hatching was stimulated by exposing eggs to light for one hour in the evening before use, and for three hours the next morning (79). To test for virus presence in each family, newly hatched coracidia were collected via centrifugation before RNA extraction.

#### Experimental infections of copepods

The first intermediate host, *Macrocyclops albidus* copepods were cultured in the laboratory at 20°C and 16:8 light:dark cycle. We used a population of copepods originating from lake Skogseidsvatnet, Norway that is highly susceptible to *S. solidus* (80). C5 copepodite stage were exposed to one coracidium each as previously described (80). Briefly, individual copepods were kept in wells of 24-well microtiter plates and starved for 2 days before exposure to a single newly hatched coracidia. Fourteen days post exposure, copepods exposed to *S. solidus* were screened under the microscope to determine the infection success. To test for the rate of vertical transmission, infected and non-infected copepods were then rinsed in sterile water, and isolated via centrifugation before RNA extraction. As controls, individual copepods that were not exposed to the parasites were collected and had their RNA extracted. To control for potential contamination by *S. solidus* in exposed but not infected copepods, we conducted PCRs with *S. solidus* specific primers according to Berger et al (81).

#### Sampling and experimental infection of threespine sticklebacks

In June 2018, we also collected mature males and gravid females of the second intermediate host *Gasterosteus aculeatus* from Rabbit slough (61° 32’ 08.1” N 149° 15’ 10.0” W), Cheney Lake, and Loberg lake and completed crosses *in vitro* to obtain lab-bred families. Fish were reared in the laboratory at 18°C and 16:8 light:dark cycle until they were 5-months old and ready for experimental exposure to *S. solidus*. Fish were fed a diet of frozen brine shrimps and chironomids larvae ad libitum daily. Each of the 224 fish were exposed to copepods parasitized with a single *S. solidus* infected with SsRV, SsJV, or neither virus. Exposure was performed by placing a single infected copepod in a tank containing a single fish that had been starved for 48h. Two days later, fish were transferred back into large tanks. After eight weeks, fish were dissected and from the five that were infected, plerocercoids, fish body cavity, liver and intestine were collected and stored in RNA later until use.

#### PCR assays for assessing viral prevalence and transmission

Specimens collected at different stages of *S. solidus* life cycle were used to assess virus prevalence, vertical transmission, and cross-species transmission to parasitized hosts. To do this, total RNA was extracted from *S. solidus* plerocercoids, culture medium used for breeding, coracidia, copepods, and stickleback tissues using the RNeasy kit (Qiagen CAT#74106) following the manufacturer’s guidelines (Qiagen CAT#74106). First strand cDNA was synthesized by reverse transcribing 500 ng of total RNA and mixed with 0.2 µg/µl Random Hexamer Primer in a 20 µl reaction volume containing 40 U/µl Ribolock™ RNase Inhibitor, 1 mM dNTPs, 200 U/µl RevertAid H Minus Reverse Transcriptase (Thermo Fisher Scientific CAT# EP0451), and water, as per manufacturer’s recommendations. Polymerase chain reaction was conducted using the Advantage 2 PCR system (Invitrogen CAT# 639137) using primers targeting the conserved RNA-dependent RNA polymerase gene of *S. solidus* associated viruses (Table S1). Amplicon presence was assayed with 1% agarose gel with SyBR Safe. Select PCR products were sequenced using Sanger sequencing to confirm primers’ specificity.

## Data mining

To assess virus presence in other populations of *S. solidus*, we queried BioProjects of publicly available transcriptomes. At the time of this study, we found PRJEB7355 (https://www.ncbi.nlm.nih.gov/bioproject/316954, 2 biosamples of wild-caught Norwegian and German *S. solidus*) which used 454 sequencing to identify plerocercoids virulence genes. BLASTn searches were used to determine the presence of 454 reads that aligned to the newly identified viruses in data from PRJEB7355. A more recent and comprehensive study, PRJNA304161 (15 biosamples from Clatworthy reservoir, England, UK) used Illumina HiSeq to compare the transcriptomes of plerocercoids collected either 70 days, 110 days or 365 days post infection of threespine sticklebacks, thus representing non-infective and infective plerocercoids, and adult stages of *S. solidus* (35). Sequence data from PRJNA304161 were downloaded and processed as follows: reads were trimmed with the Trimmomatic version 0.36 (74) with default settings. Quality filtered reads were aligned against the *S. solidus* reference genome (GCA_900618435.1) with Bowtie2 (version 2.3.4.1) (71). Unmapped reads were collected using SAMtools 1.8 (82) and bedtools (83) and assembled using the shovill method (https://github.com/tseemann/shovill). The contig sequences were compared against the GenBank non-redundant database and *S. solidus* newly discovered viruses through BLASTx. To test for virus presence in all individuals and provide relative quantitation, clean reads were aligned on assembled viruses using BWA (version 0.7.8) (84).

## Ethics statement

Stickleback collection followed guidelines for scientific fish collection by the State of Alaska Department of Fish and Game in accordance with Fish sampling permit #P17-025 and #P-18-008 and fish transport permits 17A-0024 provided to NMD. Fish were maintained at Stony Brook University under the License to collect or possess #1949 provided by the New York State Department of Environmental Conservation to NMD. Fish experiments were conducted following protocols described in Institutional Animal Care and Use Committee (IACUC) #237429 and # 815164 to Michael Bell and NMD, respectively. Fish euthanasia was conducted using MS222 and decapitation before parasite and tissue sampling. All experiments were performed in accordance with relevant guidelines and regulations in the Public Health Service Policy (PHS) on Humane Care and Use of Laboratory Animals.

## Data availability

Sequencing data were submitted to NCBI Sequence Read Archives under Bioproject accession number PRJNA576618.

## Acknowledgments

We would like to acknowledge the expertise and assistance of Dr. Michael Bell in fish collection and dissection techniques. This project was supported by the Eppley Foundation for Research and the Laurie Landeau Foundation LLC.

