## Supplementary Information for "Characterization of vertically and cross-species transmitted viruses in the cestode parasite *Schistocephalus solidus*"

Table S1. Primer pairs used to detect viruses in *Schistocephalus solidus* samples

*Primer name specifies the targeted virus. SsRV, Schistocephalus solidus rhabdovirus; SsJV, Schistocephalus solidus jingchuvirus; SsBV, Schistocephalus solidus bunya-like virus; SsTV1, Schistocephalus solidus toti-like virus 1; SsTV2, Schistocephalus solidus toti-like virus 2; SsTV3, Schistocephalus solidus toti-like virus 3.

| Primer ID | Sequence (5' – 3") | Product Length (bp) | Annealing Temp (ºC) | Purpose |
|---|---|---|---|---|
| SsRV_F3<br>SsRV_R3 | CCGTTAAGGCCGATGTTTTA<br>AGTTGACTACGCCCCAGTTG | 100 | 57 | Screening |
| SsRV_F4<br>SsRV_R4 | TTGTCAACTGGGGCGTAGTC<br>TCGTTACGGAAGGAGGAGGT | 386 | 57 | Screening |
| SsRV_F5<br>SsRV_R5 | ACCTTGTGTGGCTCGATGAT<br>GGCTGAAAATGGAAAACGAG | 500 | 52 | Bridge genome gap |
| SsRV_F6<br>SsRV_R6 | TGTGTCATTCAGGGTTTCCA<br>AGTTTGGACAGACCGCATTC | 700 | 52 | Bridge genome gap |
| SsRV_F7<br>SsRV_R7 | GCATCCTCCCTCCATCATAA<br>ACTGCAAAGTCCCAACAACC | 550 | 52 | Bridge genome gap |
| SsJV_F3<br>SsJV_R3 | TCGTCTTCCCGTAAACGAAC<br>ATTCGTACCGGACAGCACTC | 425 | 57 | Screening |
| SsJV_F4<br>SsJV_R4 | CGCTTTACCACCTTCCCTGT<br>CTTGGCGTCCGTTTCCTAGT | 280 | 57 | Screening |
| SsJV_F5<br>SsJV_R5 | TGGTGTGTGGTGTTTGGTCT<br>CCCTCGGGTAGTTCAAAGGA | 300 | 57 | Verify circular nature |
| SsBV_F1<br>SsBV_R1 | ATCATGCAGTGGACCAAGGA<br>ATGGTGTCCCTCTTGAGGTG | 900 | 55 | Screening |
| SsTV1_F1<br>SsTV1_R1 | CTCCTATACCGGTCCCCAAC<br>GCTGATAACCGCCAGAGTTC | 690 | 55 | Screening |
| SsTV2_F1<br>SsTV2_R1 | TTGGCTTTTACCAGGGTTTG<br>AAATCCAGCGTCTGACAACC | 620 | 52 | Screening |
| SsTV3_F2<br>SsTV3_R2 | CTGGAGGGGCTTAGTCTCTG<br>CAAAGCCGGAGTGATCGAAG | 970 | 55 | Screening |

Table S2. Gene specific primers (GSP) used for rapid amplification of complimentary ends (RACE) assays.

| Primer ID | Sequence (5’ – 3”) | End |
|---|---|---|
| SsRV_GSP1 | ATGTTGGCCATCTCTTTGCT | 3’ |
| SsRV_GSP2 | TCAGGAACACCTGCGTTACA | 3’ |
| SsRV_GSP3 | TGGAGAGATCGGGCAATTTA | 3’ |
| SsRV_GSP4 | CAAAGTGCCCTGGTTGTTCT | 5’ |
| SsRV_GSP5 | TGGATATAGGCGCTACATTGG | 5’ |
| SsRV_GSP6 | AGTGTTGGACTCATTGCGTC | 5’ |
| SsBV_GSP1 | GGAGAGGGAGATCGTCAACA | 3’ |
| SsBV_GSP2 | GTGGAACAGCAAACACTGGA | 3’ |
| SsBV_GSP3 | CATCACCGAGAACTTCACGA | 3’ |
| SsBV_GSP4 | GGCAAACATCACCTCCTTGT | 5’ |
| SsBV_GSP5 | AGGGCATGTTGTTGATGACA | 5’ |
| SsBV_GSP6 | ACCACCAGCAAGGTCTTCAC | 5’ |
| SsTV1_GSP1 | TCCTCAAGTCCCTGAACCAG | 3’ |
| SsTV1_GSP2 | TTACGTGGACTGAGGGCATT | 3’ |
| SsTV1_GSP3 | CACCAAAAATGTATTCGCCC | 3’ |
| SsTV1_GSP4 | GAGAAGCTGTCCACAGACG | 3’ |
| SsTV1_GSP5 | CAGTTTCTGCGTCACCCTTT | 3’ |
| SsTV1_GSP6 | TGGGCTCTCACTGTATTCGC | 3’ |
| SsTV2_GSP1 | AGTATGTCCCCGATGTGGAG | 3’ |
| SsTV2_GSP2 | CTGAGGGGACTGACTGGTGT | 3’ |
| SsTV2_GSP3 | GCTTTCCTCACAGGAACTGG | 3’ |
| SsTV2_GSP4 | GGACGAATGGATTGGAGATG | 3’ |
| SsTV3_GSP1 | CCACCCTTTCATCTGCCTAA | 3’ |
| SsTV3_GSP2 | ATGATAGGGGTGGCAGAGTG | 3’ |
| SsTV3_GSP3 | GTTTGAAGCCATGGGAGAAC | 3’ |
| SsTV3_GSP4 | ACCTGGCAGAGGCAATTAGA | 3’ |
| SsTV3_GSP5 | AAAGATGAATGGGTCGGTGA | 3’ |

| | | |
|---|---|---|
| SsTV3_GSP6 | TCTATTCGGGCCTACAGGAG | 3’ |

Table S3: transcription initiation and termination signals in SsRV genome

| **ORF** | **Intergenic** | **Transcription initiation sequence** | **Termination signal** |
|---|---|---|---|
| N | | **UUGU**UGUGUAUAUUUGCUUUUAACAGUGAACGCCCCAUAC | UCUUUUUUU |
| P | G | **UUGU**UAC | UCUUUUUUU |
| M | G | **UUGU**UUUUUCCUUUUUGGCUCUGUAGUAC | UCUUUUUUU |
| G | G | **UUGU**GUUGUGGAACUUGGGAUGUCGUUCUUGGUCUUAAGGUAACGCUUGCACUCUUAC | UCUUUUUUU |
| α1 | G | **UUGU**AGAUUGACGGGGCCCCUCCUGUGUGUAC | UCUUUUUUU |
| L | G | **UUGU**UAGUAAACCAUUCGAUCUGUUUAACAAUGACACUAACUAGUAC | UCUUUUUUU |

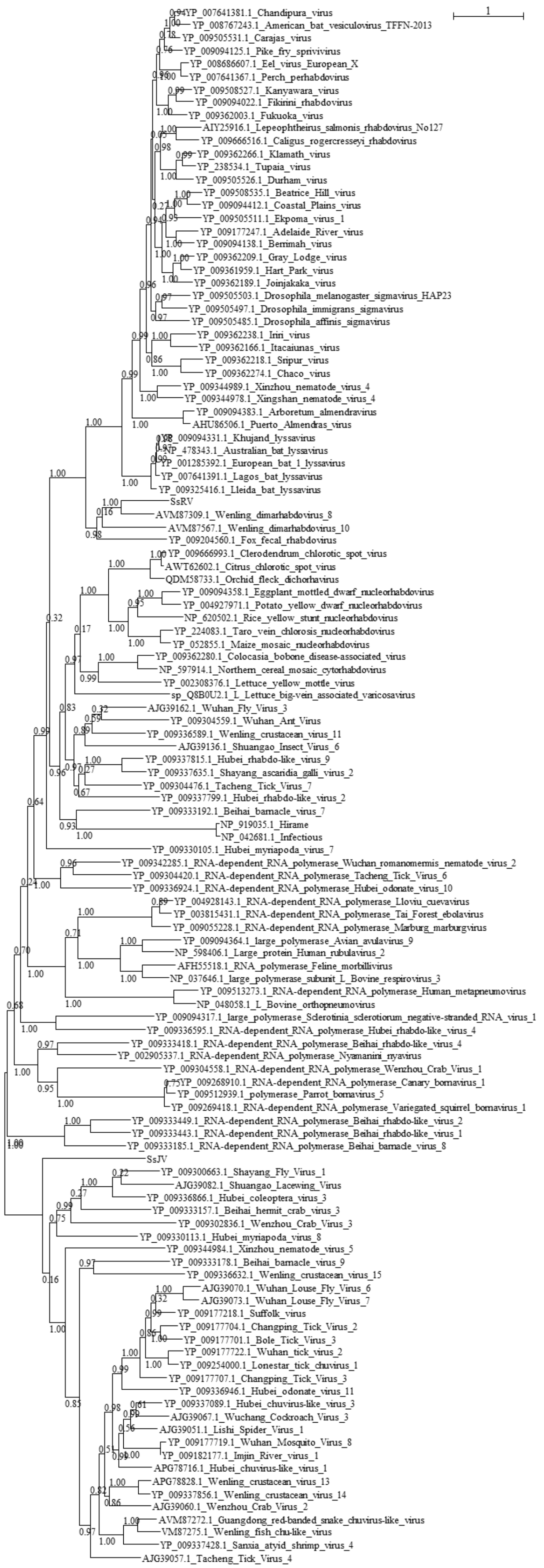
1
YP_007641381.1_Chandipura_virus
YP_008767243.1_American_bat_vesiculovirus_TFFN-2013
YP_009505531.1_Carajas_virus
YP_009094125.1_Pike_fry_sprivivirus
YP_008686607.1_Eel_virus_European_X
YP_007641367.1_Perch_perhabdovirus
YP_009508527.1_Kanyawara_virus
YP_009094022.1_Fikirini_rhabdovirus
YP_009362003.1_Fukuoka_virus
AIY25916.1_Lepeophtheirus_salmonis_rhabdovirus_No127
YP_009666516.1_Caligus_rogercresseyi_rhabdovirus
YP_009362266.1_Klamath_virus
YP_238534.1_Tupaia_virus
YP_009505526.1_Durham_virus
YP_009508535.1_Beatrice_Hill_virus
YP_009094412.1_Coastal_Plains_virus
YP_009505511.1_Ekpoma_virus_1
YP_009177247.1_Adelaide_River_virus
YP_009094138.1_Berrimah_virus
YP_009362209.1_Gray_Lodge_virus
YP_009361959.1_Hart_Park_virus
YP_009362189.1_Joinjakaka_virus
YP_009505503.1_Drosophila_melanogaster_sigmavirus_HAP23
YP_009505497.1_Drosophila_immigrans_sigmavirus
YP_009505485.1_Drosophila_affinis_sigmavirus
YP_009362238.1_Iriri_virus
YP_009362166.1_Itacaiunas_virus
YP_009362218.1_Sripur_virus
YP_009362274.1_Chaco_virus
YP_009344989.1_Xinzhou_nematode_virus_4
YP_009344978.1_Xingshan_nematode_virus_4
YP_009094383.1_Arboretum_almendravirus
AHU86506.1_Puerto_Almendras_virus
YP_009094331.1_Khujand_lyssavirus
NP_478343.1_Australian_bat_lyssavirus
YP_001285392.1_European_bat_1_lyssavirus
YP_007641391.1_Lagos_bat_lyssavirus
YP_009325416.1_Lleida_bat_lyssavirus
SsRV
AVM87309.1_Wenling_dimarhabdovirus_8
AVM87567.1_Wenling_dimarhabdovirus_10
YP_009204560.1_Fox_fecal_rhabdovirus
YP_009666993.1_Clerodendrum_chlorotic_spot_virus
AWT62602.1_Citrus_chlorotic_spot_virus
QDM58733.1_Orchid_fleck_dichorhavirus
YP_009094358.1_Eggplant_mottled_dwarf_nucleorhabdovirus
YP_004927971.1_Potato_yellow_dwarf_nucleorhabdovirus
NP_620502.1_Rice_yellow_stunt_nucleorhabdovirus
YP_224083.1_Taro_vein_chlorosis_nucleorhabdovirus
YP_052855.1_Maize_mosaic_nucleorhabdovirus
YP_009362280.1_Colocasia_bobone_disease-associated_virus
NP_597914.1_Northern_cereal_mosaic_cytorhabdovirus
YP_002308376.1_Lettuce_yellow_mottle_virus
sp_Q8B0U2.1_L_Lettuce_big-vein_associated_varicosavirus
AJG39162.1_Wuhan_Fly_Virus_3
YP_009304559.1_Wuhan_Ant_Virus
YP_009336589.1_Wenling_crustacean_virus_11
AJG39136.1_Shuangao_Insect_Virus_6
YP_009337815.1_Hubei_rhabdo-like_virus_9
YP_009337635.1_Shayang_ascaridia_galli_virus_2
YP_009304476.1_Tacheng_Tick_Virus_7
YP_009337799.1_Hubei_rhabdo-like_virus_2
YP_009333192.1_Beihai_barnacle_virus_7
NP_919035.1_Hirame
NP_042681.1_Infectious
YP_009330105.1_Hubei_myriapoda_virus_7
YP_009342285.1_RNA-dependent_RNA_polymerase_Wuchan_romanomermis_nematode_virus_2
YP_009304420.1_RNA-dependent_RNA_polymerase_Tacheng_Tick_Virus_6
YP_009336924.1_RNA-dependent_RNA_polymerase_Hubei_odonate_virus_10
YP_004928143.1_RNA-dependent_RNA_polymerase_Lloviu_cuevavirus
YP_003815431.1_RNA-dependent_RNA_polymerase_Tai_Forest_ebolavirus
YP_009055228.1_RNA-dependent_RNA_polymerase_Marburg_marburgvirus
YP_009094364.1_large_polymerase_Avian_avulavirus_9
NP_598406.1_Large_protein_Human_rubulavirus_2
AFH55518.1_RNA_polymerase_Feline_morbillivirus
NP_037646.1_large_polymerase_subunit_L_Bovine_respirovirus_3
YP_009513273.1_RNA-dependent_RNA_polymerase_Human_metapneumovirus
NP_048058.1_L_Bovine_orthopneumovirus
YP_009094317.1_large_polymerase_Sclerotinia_sclerotiorum_negative-stranded_RNA_virus_1
YP_009336595.1_RNA-dependent_RNA_polymerase_Hubei_rhabdo-like_virus_4
YP_009333418.1_RNA-dependent_RNA_polymerase_Beihai_rhabdo-like_virus_4
YP_002905337.1_RNA-dependent_RNA_polymerase_Nyamanini_nyavirus
YP_009304558.1_RNA-dependent_RNA_polymerase_Wenzhou_Crab_Virus_1
YP_009268910.1_RNA-dependent_RNA_polymerase_Canary_bornavirus_1
YP_009512939.1_polymerase_Parrot_bornavirus_5
YP_009269418.1_RNA-dependent_RNA_polymerase_Variegated_squirrel_bornavirus_1
YP_009333449.1_RNA-dependent_RNA_polymerase_Beihai_rhabdo-like_virus_2
YP_009333443.1_RNA-dependent_RNA_polymerase_Beihai_rhabdo-like_virus_1
YP_009333185.1_RNA-dependent_RNA_polymerase_Beihai_barnacle_virus_8
SsJV
YP_009300663.1_Shayang_Fly_Virus_1
AJG39082.1_Shuangao_Lacewing_Virus
YP_009336866.1_Hubei_coleoptera_virus_3
YP_009333157.1_Beihai_hermit_crab_virus_3
YP_009302836.1_Wenzhou_Crab_Virus_3
YP_009330113.1_Hubei_myriapoda_virus_8
YP_009344984.1_Xinzhou_nematode_virus_5
YP_009333178.1_Beihai_barnacle_virus_9
YP_009336632.1_Wenling_crustacean_virus_15
AJG39070.1_Wuhan_Louse_Fly_Virus_6
AJG39073.1_Wuhan_Louse_Fly_Virus_7
YP_009177218.1_Suffolk_virus
YP_009177704.1_Changping_Tick_Virus_2
YP_009177701.1_Bole_Tick_Virus_3
YP_009177722.1_Wuhan_tick_virus_2
YP_009254000.1_Lonestar_tick_chuvirus_1
YP_009177707.1_Changping_Tick_Virus_3
YP_009336946.1_Hubei_odonate_virus_11
YP_009337089.1_Hubei_chuvirus-like_virus_3
AJG39067.1_Wuchang_Cockroach_Virus_3
AJG39051.1_Lishi_Spider_Virus_1
YP_009177719.1_Wuhan_Mosquito_Virus_8
YP_009182177.1_Imjin_River_virus_1
APG78716.1_Hubei_chuvirus-like_virus_1
APG78828.1_Wenling_crustacean_virus_13
YP_009337856.1_Wenling_crustacean_virus_14
AJG39060.1_Wenzhou_Crab_Virus_2
AVM87272.1_Guangdong_red-banded_snake_chuvirus-like_virus
VM87275.1_Wenling_fish_chu-like_virus
YP_009337428.1_Sanxia_atyid_shrimp_virus_4
AJG39057.1_Tacheng_Tick_Virus_4

Figure S1: Phylogenetic analysis of the RdRP of viruses from the orders *Mononegavirales* and *Jingchuvirales*.

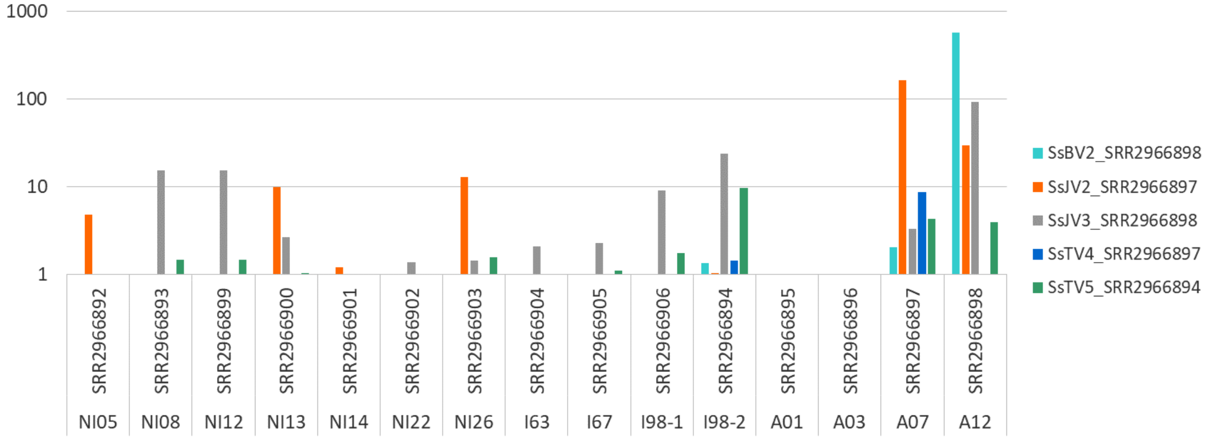


Figure S2: Virus prevalence in samples from PRJNA304161. The figure shows the number of reads mapped against the five viruses assembled from this dataset (SsJV2, SsJV3, SsBV2, SsTV4 and SsTV5) per millions of clean reads for each of the 15 individual parasites. NI: non-infective plerocercoids; I: infective plerocercoids; A: adults.

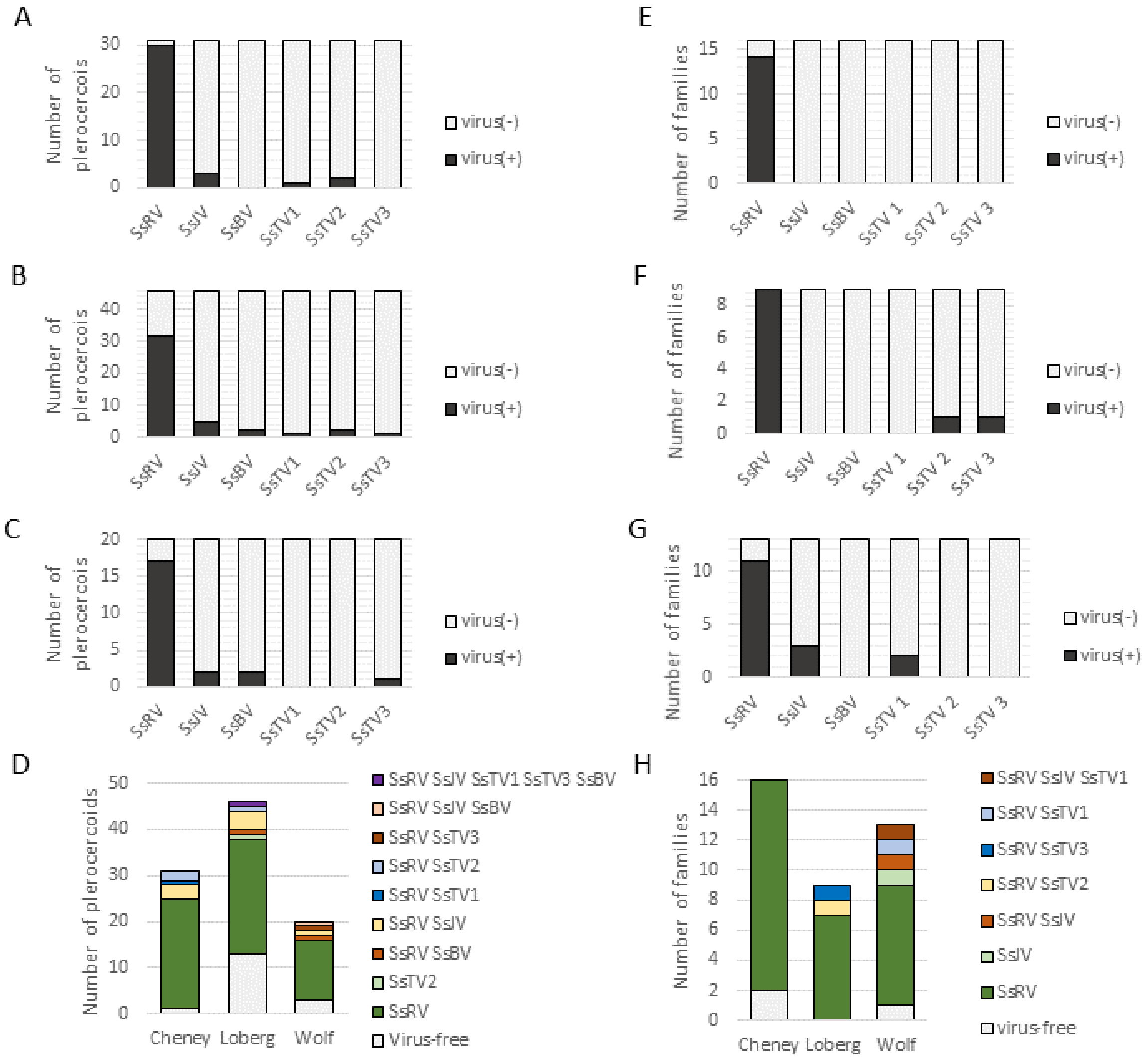


Figure S3: Virus prevalence in plerocercoids (A-D) and presence in families from *in vitro* breeding (E-F). Virus presence in Cheney lake (A and E), Loberg Lake (B and F), and Wolf Lake (C and G). Plerocercoids and families were often coinfected by multiple viruses (D and H).

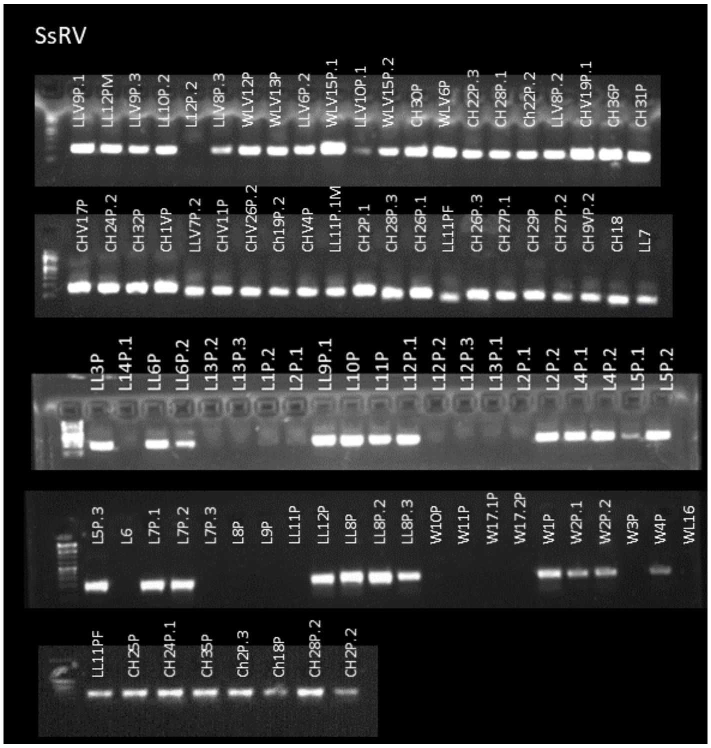


Figure S4: Detection of SsRV in plerocercoids from Cheney Lake (CL, 31 individuals), Loberg Lake (LL, 46 individuals) and Wolf lake (WL, 20 individuals). .1 .2 and .3 is used to label plerocercoids collected from the same stickleback host.

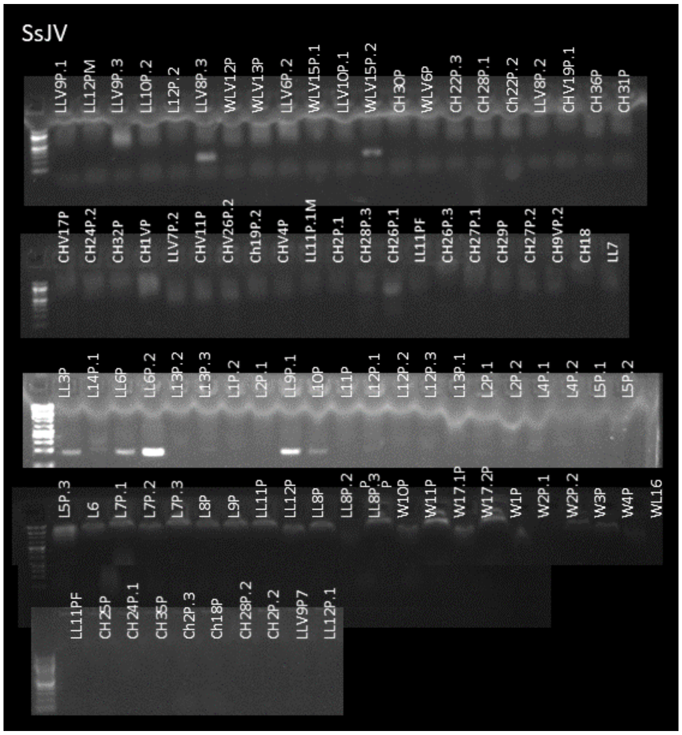

Figure S5: Detection of SsJV in plerocercoids from Cheney Lake (CL, 31 individuals), Loberg Lake (LL, 46 individuals) and Wolf lake (WL, 20 individuals). .1 .2 and .3 is used to label plerocercoids collected from the same stickleback host.

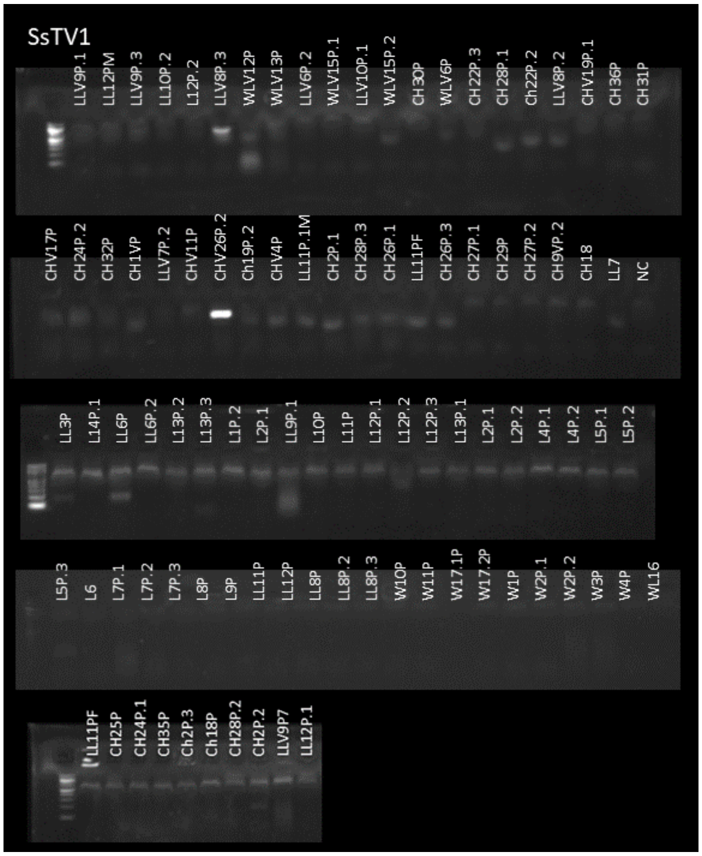


Figure S6: Detection of SsTV1 in plerocercoids from Cheney Lake (CL, 31 individuals), Loberg Lake (LL, 46 individuals) and Wolf lake (WL, 20 individuals). .1 .2 and .3 is used to label plerocercoids collected from the same stickleback host.

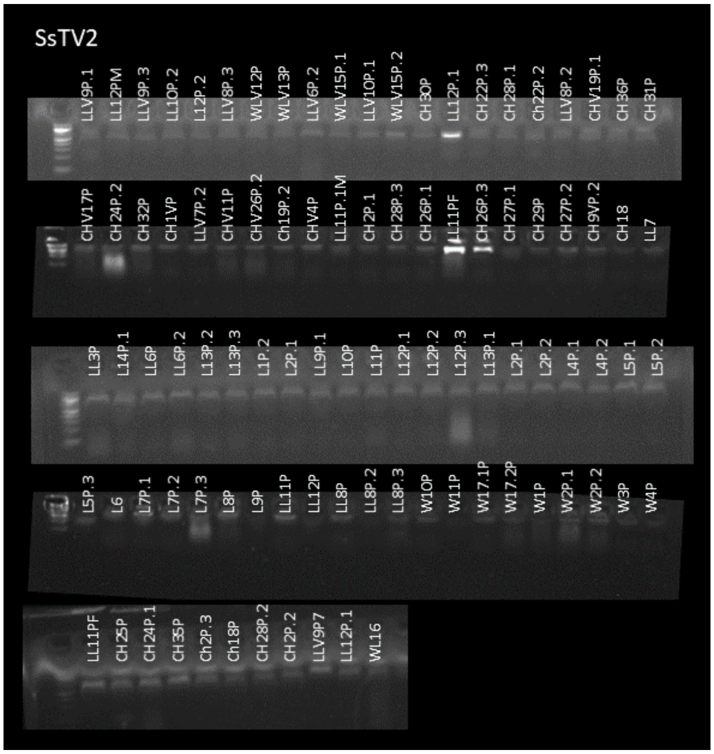


Figure S7: Detection of SsTV2 in plerocercoids from Cheney Lake (CL, 31 individuals), Loberg Lake (LL, 46 individuals) and Wolf lake (WL, 20 individuals). .1 .2 and .3 is used to label plerocercoids collected from the same stickleback host.

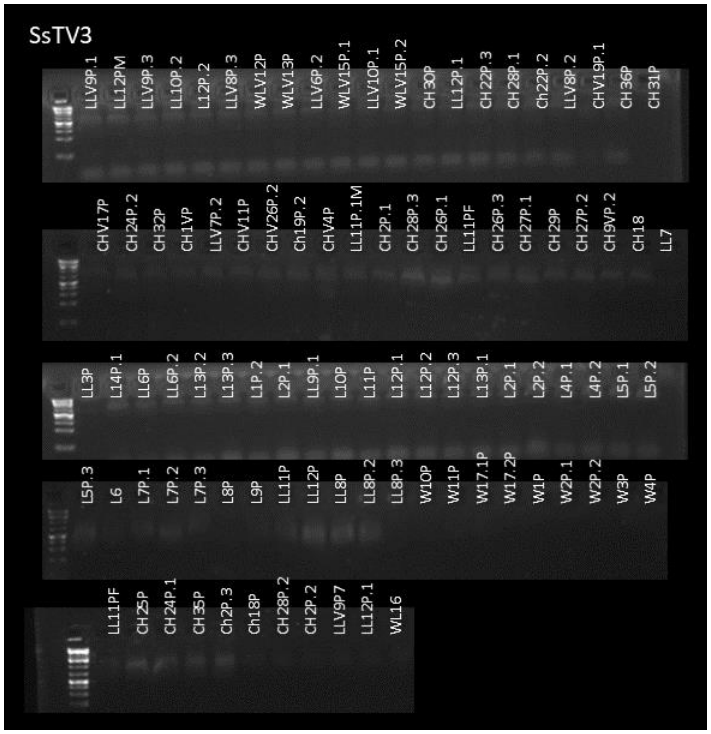

Figure S8: Detection of SsTV3 in plerocercoids from Cheney Lake (CL, 31 individuals), Loberg Lake (LL, 46 individuals) and Wolf lake (WL, 20 individuals). .1 .2 and .3 is used to label plerocercoids collected from the same stickleback host.

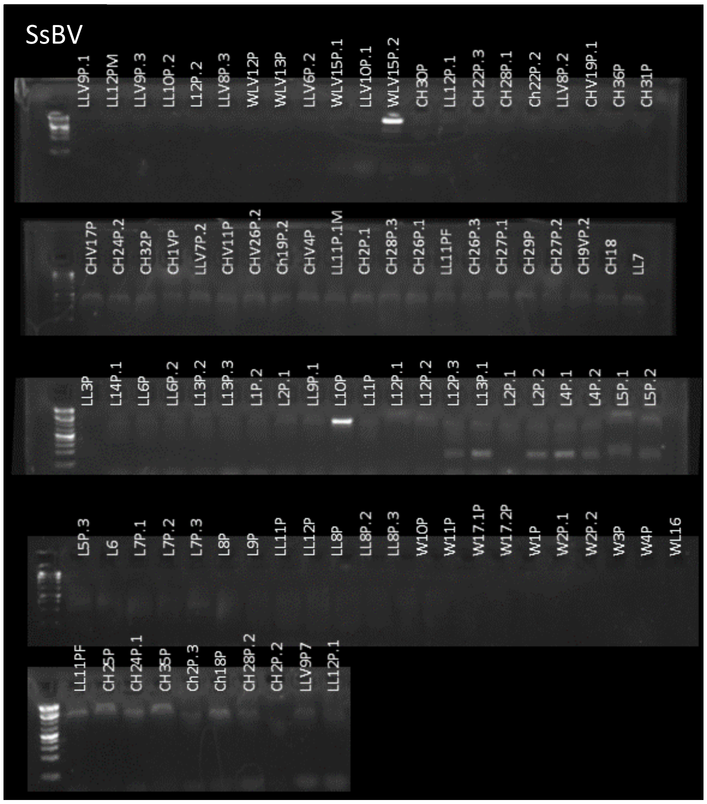


Figure S9: Detection of SsBV in plerocercoids from Cheney Lake (CL, 31 individuals), Loberg Lake (LL, 46 individuals) and Wolf lake (WL, 20 individuals). .1 .2 and .3 is used to label plerocercoids collected from the same stickleback host.

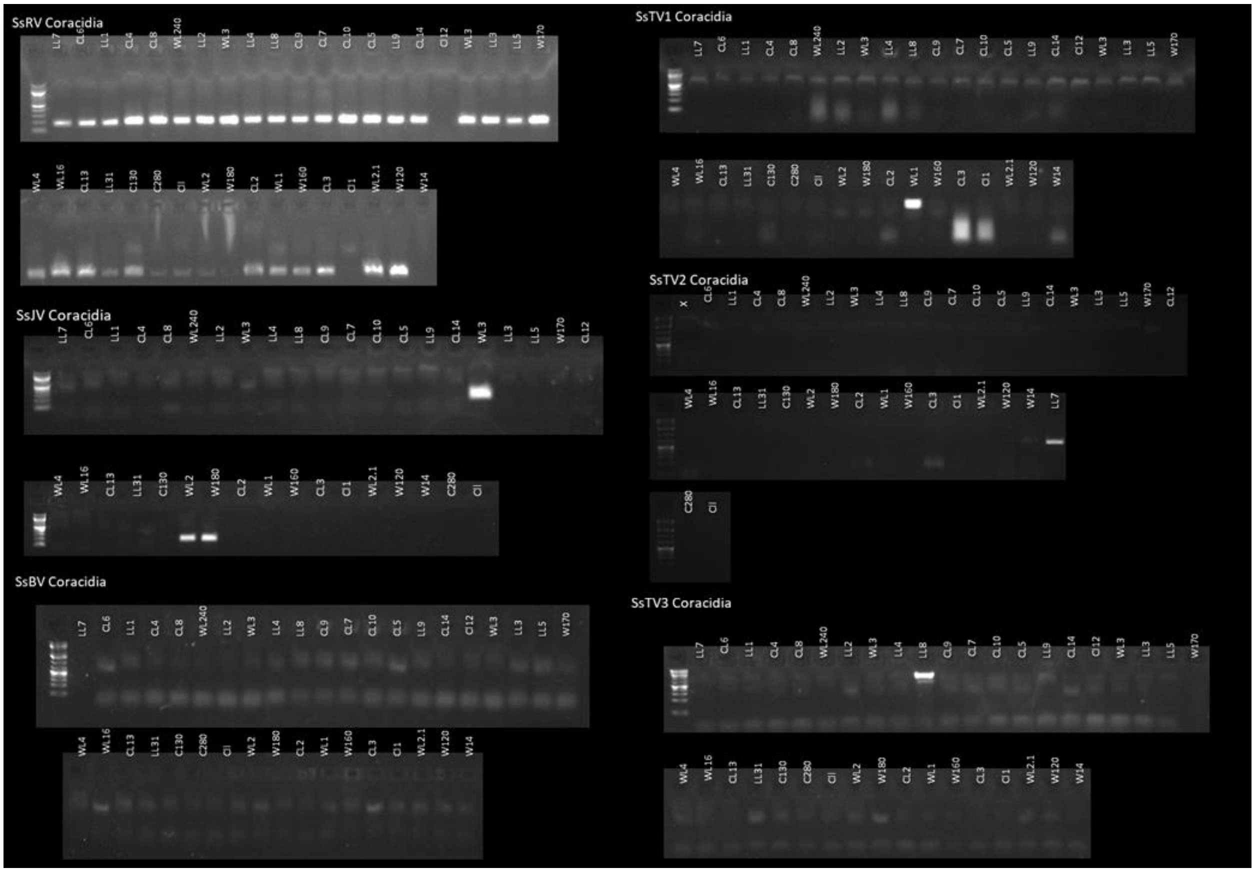


Figure S10: Detection of SsRV, SsJV, SsBV, SsTV1, SsTV2 and SsTV3 in coracidia from 38 families produced *in vitro* from plerocercoids collected in Cheney Lake (CL, 16 families), Loberg Lake (LL, 9 families) and Wolf lake (WL, 13 families).

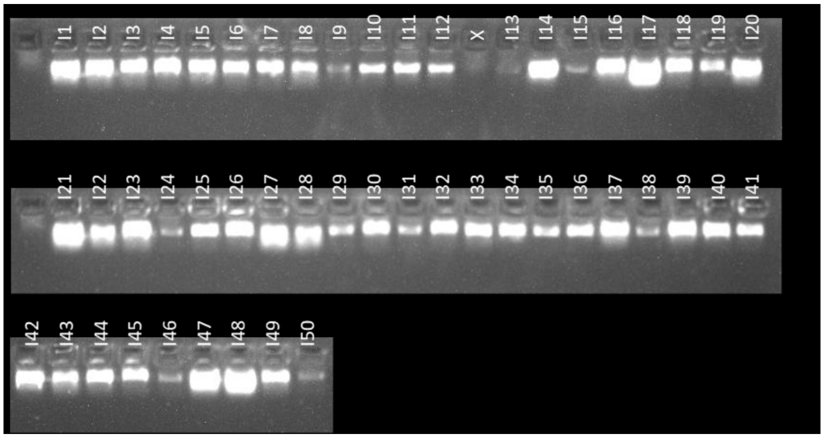


Figures S11: Detection of SsRV in procercoids following experimental infection of copepods with coracidia from the SsRV(+) family WL1.

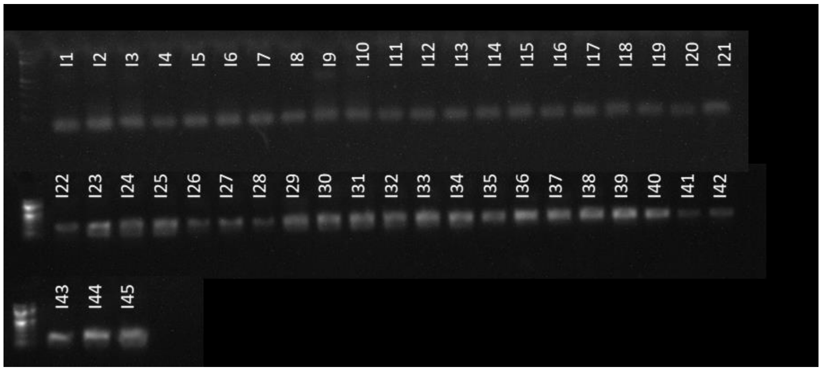


Figures S12: Detection of SsJV in procercoids following experimental infection of copepods with coracidia from the SsJV(+) family WL240.

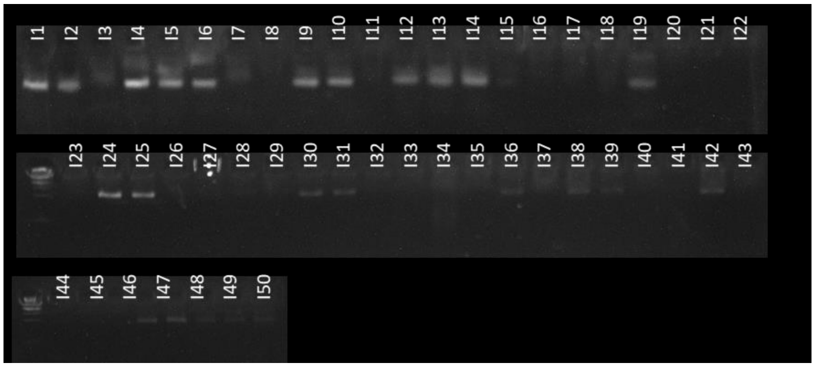


Figure S13: Detection of SsTV1 in procercoids following experimental infection of copepods with coracidia from the SsTV1(+) family WL1.

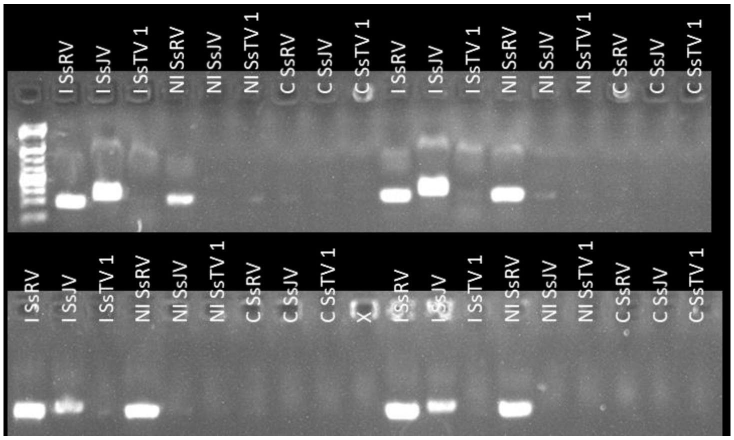


Figure S14: Detection of SsRV, SsJV, and SsTV1 in copepods experimentally infected by *S. solidus*. C: control non-exposed NI: exposed but non-infected I: infected.

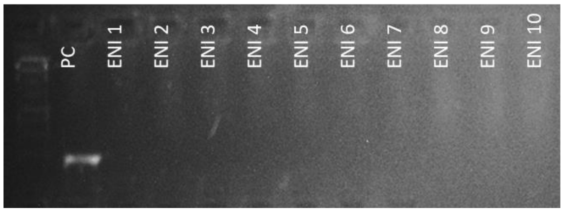


Figure S15: Detection of *S. solidus* in copepods exposed to *S. solidus* but not infected (ENI) and a positive control (PC).

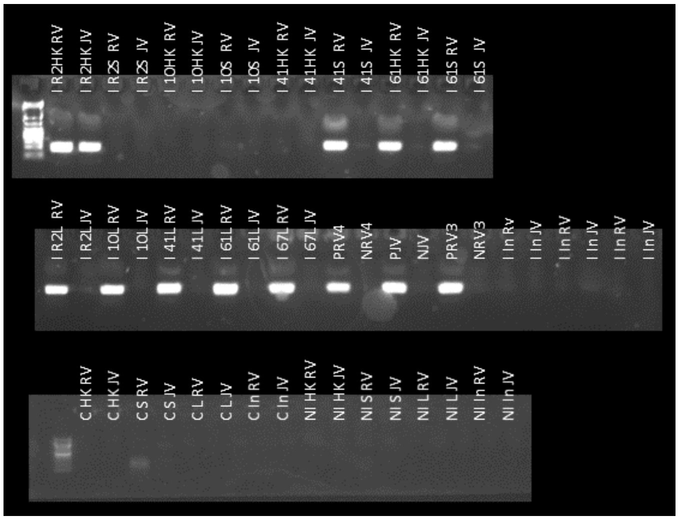


Figure S16: Detection of SsRV and SsJV in head kidney (HK), spleen (S), Liver (L), and intestine (In) tissues of Threespine sticklebacks experimentally infected by *S. solidus*. C: control non-exposed NI: exposed but non-infected I: infected. RV: SsRV; JV: SsJV

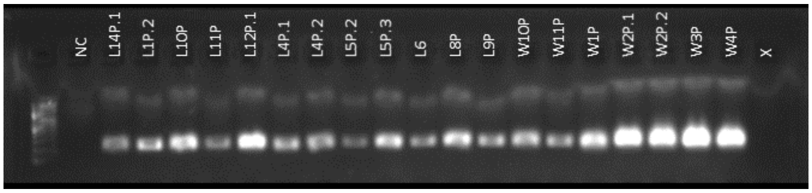


Figure S17: Detection of SsRV in liver tissue from field collected sticklebacks. The presence of SsRV in plerocercoids from corresponding fish individuals is provided in Figure 1. NC represents the negative control.

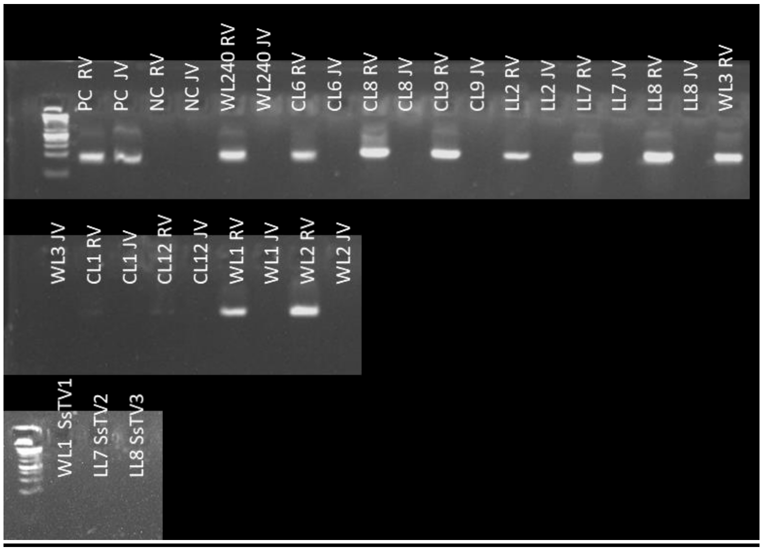


Figure S18: Detection of SsRV, SsJV, and SsTV1 in the culture medium used for in vitro breeding of S. solidus families. Presence of the corresponding viruses in coracidia of corresponding families is available in supplementary figure 2. Positive control (PC) and negative control (NC) were also included.
